# Normothermic Ex-vivo Kidney Perfusion in a Porcine Auto-Transplantation Model Preserves the Expression of Key Mitochondrial Proteins: An Unbiased Proteomics Analysis

**DOI:** 10.1101/2020.08.17.253252

**Authors:** Caitriona M. McEvoy, Sergi Clotet-Freixas, Tomas Tokar, Chiara Pastrello, Shelby Reid, Ihor Batruch, Adrien A.E. RaoPeters, J. Moritz Kaths, Peter Urbanellis, Sofia Farkona, Julie A.D. Van, Bradley L. Urquhart, Rohan John, Igor Jurisica, Lisa A. Robinson, Markus Selzner, Ana Konvalinka

## Abstract

Normothermic *ex-vivo* kidney perfusion (NEVKP) results in significantly improved graft function in porcine auto-transplant models of DCD injury compared to static cold storage (SCS); however, the molecular mechanisms underlying these beneficial effects remain unclear. We performed an unbiased proteomics analysis of 28 kidney biopsies obtained at 3 time points from pig kidneys subjected to 30-minutes of warm ischemia, followed by 8 hours of NEVKP or SCS, and auto-transplantation. 70/6593 proteins quantified were differentially expressed between NEVKP and SCS groups (FDR<0.05). Proteins increased in NEVKP mediated key metabolic processes including fatty acid ß-oxidation, the TCA-cycle and oxidative phosphorylation. Comparison of our findings with external datasets of ischemia-reperfusion, and other models of kidney injury confirmed that 47 of our proteins represent a common signature of kidney injury reversed or attenuated by NEVKP. We validated key metabolic proteins (ETFB, CPT2) by immunoblotting. Transcription factor databases identified PPARGC1A, PPARA/G/D and RXRA/B as the upstream regulators of our dataset, and we confirmed their increased expression in NEVKP with RT-PCR. The proteome-level changes observed in NEVKP mediate critical metabolic pathways that may explain the improved graft function observed. These effects may be coordinated by PPAR-family transcription factors, and may represent novel therapeutic targets in ischemia-reperfusion injury.

## Introduction

Kidney transplantation is considered the optimal treatment for patients with end-stage kidney disease (ESKD).^1–4^ The increased prevalence of ESKD in recent years has led to a growing demand for renal transplantation^5,6^ which exceeds the organ supply.^7,8^ Increased utilization of marginal grafts, i.e. from donation after circulatory death (DCD) and extended criteria donors is incentivized in the face of organ shortage.^7,9,10^ While these organs confer a survival benefit in comparison to remaining on dialysis,^10^ studies have demonstrated inferior allograft outcomes compared to standard criteria donor grafts; including increased rates of primary non-function, delayed graft function (DGF), and less favorable graft outcomes at one year.^11–17^ Prolonged cold ischemic time, and warm ischemic time-characteristic of DCD, are significant risk factors for these adverse outcomes. DCD kidneys, particularly, are poorly tolerant of cold ischemia, and more susceptible to ischemia-reperfusion injury (IRI).^15–19^ The increased utilization of DCD kidneys renewed focus on optimizing organ preservation; particularly on machine perfusion alternatives to the cold anoxic storage methods (static cold storage (SCS) and hypothermic machine perfusion) currently in widespread use.^20^ Normothermic *ex-vivo* kidney perfusion (NEVKP) shows particular promise. While cold anoxic storage is associated with suspended cell metabolism, NEVKP provides a continuous flow of warmed, oxygenated perfusate containing nutritional substrates, thereby maintaining the metabolic activity of the tissue in a near-physiologic state.^21,22^ Consequently, NEVKP permits graft assessment, conditioning, and repair throughout perfusion.^23^ NEVKP results in superior short-term outcomes when compared with SCS in a porcine DCD auto-transplantation model.^21,24–28^ Assessment of perfusion characteristics and biomarkers during NEVKP allowed prediction of post-transplant graft function,^29^ highlighting the potential of NEVKP to inform decision-making regarding organ suitability for transplantation. Normothermic perfusion is successfully applied in other solid-organ transplant settings.^30–32^ In kidney transplantation, the first clinical trial of short (1hour) NEVKP after hypothermic preservation showed positive results,^33^ with further studies ongoing.

Despite the observed benefits, the molecular mechanisms responsible for improved graft function with NEVKP remain undefined. We hypothesized that NEVKP would induce key alterations in the renal proteome compared to SCS in a DCD model, and that identifying these changes would provide insights into the molecular mechanisms central to superior graft function in this setting.

## Methods

### Study Design

We conducted an unbiased proteomics analysis in a porcine DCD auto-transplantation model comprising two groups (8 hours NEVKP and 8 hours SCS), n=5 animals/group. Kidney biopsy tissue was collected at three timepoints: baseline (contralateral kidney, prior to warm ischemia), 30minutes post-reperfusion, and at sacrifice (post-operative day 3 (POD3)) (Figure 1a). All samples were snap-frozen in liquid nitrogen, and stored at −80°C.

**Figure 1:**
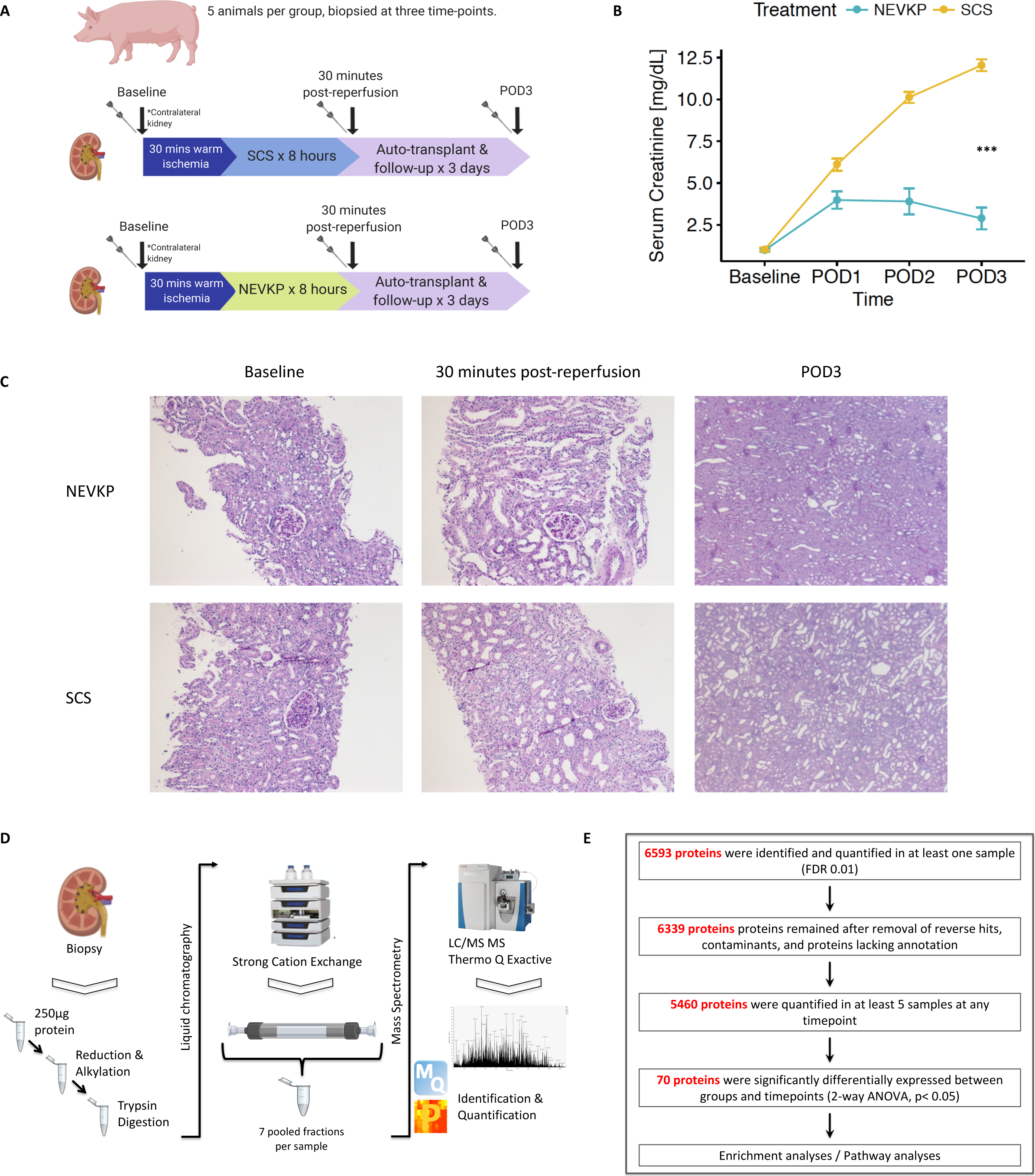
Overview of experimental model and proteomics workflow. (**A**) Details of porcine DCD auto-transplantation model comprising of two groups (8 hours NEVKP and 8 hours SCS), n=5 animals/group; biopsied at three timepoints: baseline (from the contralateral kidney, prior to warm ischemia), 30 minutes post-reperfusion, and at sacrifice (POD3). (**B**) Interaction plot showing serum creatinine (mean ± SEM in mg/dL) of the transplanted animals during 3-day post-operative follow-up in NEVKP-and SCS-treated groups respectively (Data amended from reference 34). A polynomial regression of creatinine levels in dependence on treatment, time and time2 was performed (F-test, p-value < 2.23x 10^-15^). (**C**) Light microscopy of PAS-stained images from representative NEVKP-treated (top panel) and SCS-treated (bottom panel) kidneys. Images from baseline (10X), 30minutes post-reperfusion (10X) and post-operative day 3 (POD3) (2.5X) are shown. (**D**) Simplified proteomics workflow including sample processing, strong cation exchange liquid chromatography and fractionation, followed by LC-MS/MS on a Thermo Q Exactive Plus mass spectrometer, and subsequent identification and quantification of peptides is shown. (**E**) Overview of proteomics data analysis including the numbers of identified and quantified proteins, and the number of proteins differentially expressed between groups and across timepoints (2-way ANOVA with Tukey’s HSD correction). Proteins with q-value <0.05 for the effect of treatment and time were considered differentially expressed. LC-MS/MS, liquid chromatography followed by tandem mass spectrometry; NEVKP, Normothermic ex vivo kidney perfusion; PAS, periodic acid Schiff; POD3, post-operative day 3; SCS, static cold storage.

### Experimental model and NEVKP

Following induction of general anesthesia, the right renal artery and vein were clamped for 30 minutes, mimicking a DCD-type injury, as described previously.^24,27^ The right kidney was then removed, flushed, and subjected to 8 hours of either SCS or NEVKP prior to re-implantation^24,27^. Animals were followed-up for 3 days before euthanization.

### Proteomics sample preparation

Frozen porcine kidney biopsy samples were lysed, homogenized and sonicated. The supernatant was collected following centrifugation at 15,000g at 4°C for 20 minutes. Total protein concentration was measured using Coomassie assay and each sample was normalized to 250μg of total protein. Two samples had <100μg of total protein and were eliminated. The remaining samples underwent denaturation, reduction and alkylation before trypsinization overnight at 37°C. SCX chromatography and peptide fractionation were performed on an HPLC system (Agilent1100) using a 60-minute two-step gradient. Eluting peptides were pooled into 7 fractions.

### Tandem mass spectrometry

Peptides were identified by LC-MS/MS as described previously^34^. Briefly, peptides from each fraction were eluted and subjected to liquid chromatography coupled online to a Q-Exactive Plus mass spectrometer. The top 12 peaks were selected for MS/MS. For protein identification and data analysis, XCalibur software (ThermoFisher) was utilized to generate RAW files of each MS run.

### Proteomic Data analysis

The raw mass spectra from each fraction were analysed using Andromeda search engine (MaxQuant(v.1.5.3.28)) against a Sus scrofa database generated from the non-redundant union of porcine sequences from UniProtKB, NCBI-RefSeq, and cRAP database of common contaminants^35^. Reverse decoy mode was used. Tryptic peptides were selected with up to two mis-cleavages. Methionine oxidation and N-terminal protein acetylation were selected as variable modifications. Carbamidomethylation was selected as fixed modification.

Protein/site FDR were set at 0.01. Label-free quantification was performed and normalized protein LFQ intensities were calculated.

The data were analysed using Perseus (v.1.5.2.6). Reverse hits and contaminants were removed. Normalized LFQ intensities were log2-transformed. The data were filtered to only retain proteins identified in at least 5 samples at any timepoint. The MS data have been deposited to the ProteomeXchange Consortium via the PRIDE partner repository^36^ with the dataset identifier PXD015277.

The protein expression data were prepossessed by imputing the missing values using the QRILC method from the R package imputeLCMD(v2.0) under default parametrization^37^. 300 independent replicates of the imputed data were created. For each, we performed 2-way ANOVA followed by Tukey’s HSD test. The p-values were adjusted for multiple testing by the FDR method. Robust estimates of statistical significance were calculated using geometric mean of the adjusted p-values across the imputation replicates.

For dataset clustering and visualization, we used Binary Tree-Structured Vector Quantization (BTSVQ) algorithm^38^. Gene Ontology and pathway enrichment were calculated using g:Profiler^39^ and pathDIP^40^ respectively. Upstream regulator analysis was performed using ARCHS4^41^ and Catrin (http://142.150.188.233:9080/Catrin/index.jsp).

### Urine Metabolomics

Indoxyl sulfate, p-cresyl sulfate, p-cresyl glucuronide, hippuric acid, betaine and choline were quantified using Ultra Performance Liquid Chromatography coupled to Quadrupole Time-of-Flight mass spectrometry. Data were acquired in sensitivity mode with a 0.05 second scan-time in a 50-1200 *m/z* range and the *m/z* of each analyte was specifically targeted. Analytes were quantified using TargetLynx V4.1 software (Waters) by comparing sample peaks to a twelve-point standard curve generated for each metabolite. Quality control samples containing known concentrations of each analyte were prepared using the same protocol and injected every nine samples. The coefficient of variation of the assay was < 10% for all analytes.

### Statistical Analysis

Significance between groups was assessed by Mann-Whitney test. P*-*values <0.05 were considered significant. Urinary metabolite concentrations were adjusted for urinary creatinine concentration. POD3 values were expressed as fold change over baseline. R (v3.5.2) and GraphPad Prism software (v8) were used for analysis and graph preparation. *p<0.05, **p<0.01, ***p<0.001.

**The supplement includes a detailed description of all methods.**

## Results

### Proteomic analysis of NEVKP and SCS biopsies

There were 5 animals per experimental group, each biopsied at 3 timepoints (n=30 biopsies) (Figure 1A). Two biopsies with insufficient protein yield (<100μg) to generate comparable results to the remaining biopsies were excluded. As previously reported by our group,^42^ NEVKP-preserved grafts demonstrated superior kidney function after heterotopic auto-transplantation compared to SCS-preserved grafts, with significantly lower serum creatinine (SCr) post-operatively in the NEVKP group compared to the SCS (Figure 1B)(F-test, p<2.23×10^-15^). Light microscopy demonstrated normal histology at baseline, with mild tubular injury in both groups at 30minutes post-reperfusion, slightly more prominent in SCS (Figure 1C), as previously reported in this model.^42^ Tubular injury and dilatation was evident at POD3, and was more severe in SCS-treated kidneys (Figure 1C).

28 samples comprising 9 baseline samples (4 NEVKP, 5 SCS), 9 samples from 30minutes post-reperfusion (4 NEVKP, 5 SCS), and 10 samples from POD3 (5 NEVKP, 5 SCS) were analyzed by LC-MS/MS, as summarized in Figure 1D.

6593 proteins were identified and quantified in ≥1 sample (FDR<0.01) (Figure 1E). After removal of contaminants, reverse hits and proteins lacking annotation, 6339 proteins remained. Of these, 5460 proteins were quantified in ≥5 samples at any timepoint, and remained in the final dataset for analysis. Missing values were then imputed and, as expected, represented the low abundance proteins (**Supplemental Figure 1**). 70 proteins were identified as differentially expressed between experimental groups and timepoints (2-way ANOVA with Tukey’s HSD post-hoc test, adjusted p-value<0.05) (Table 1).

**Table 1:**
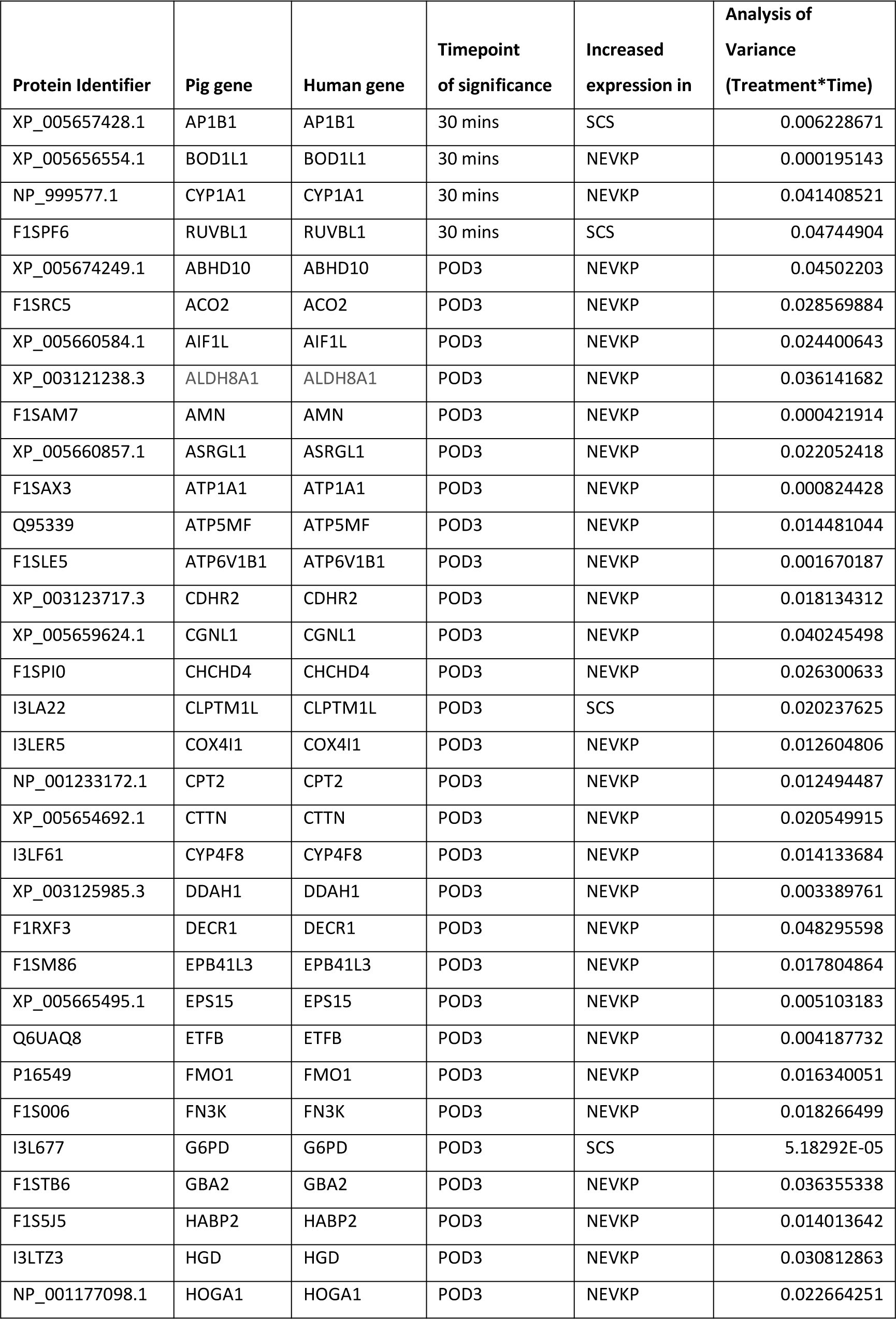
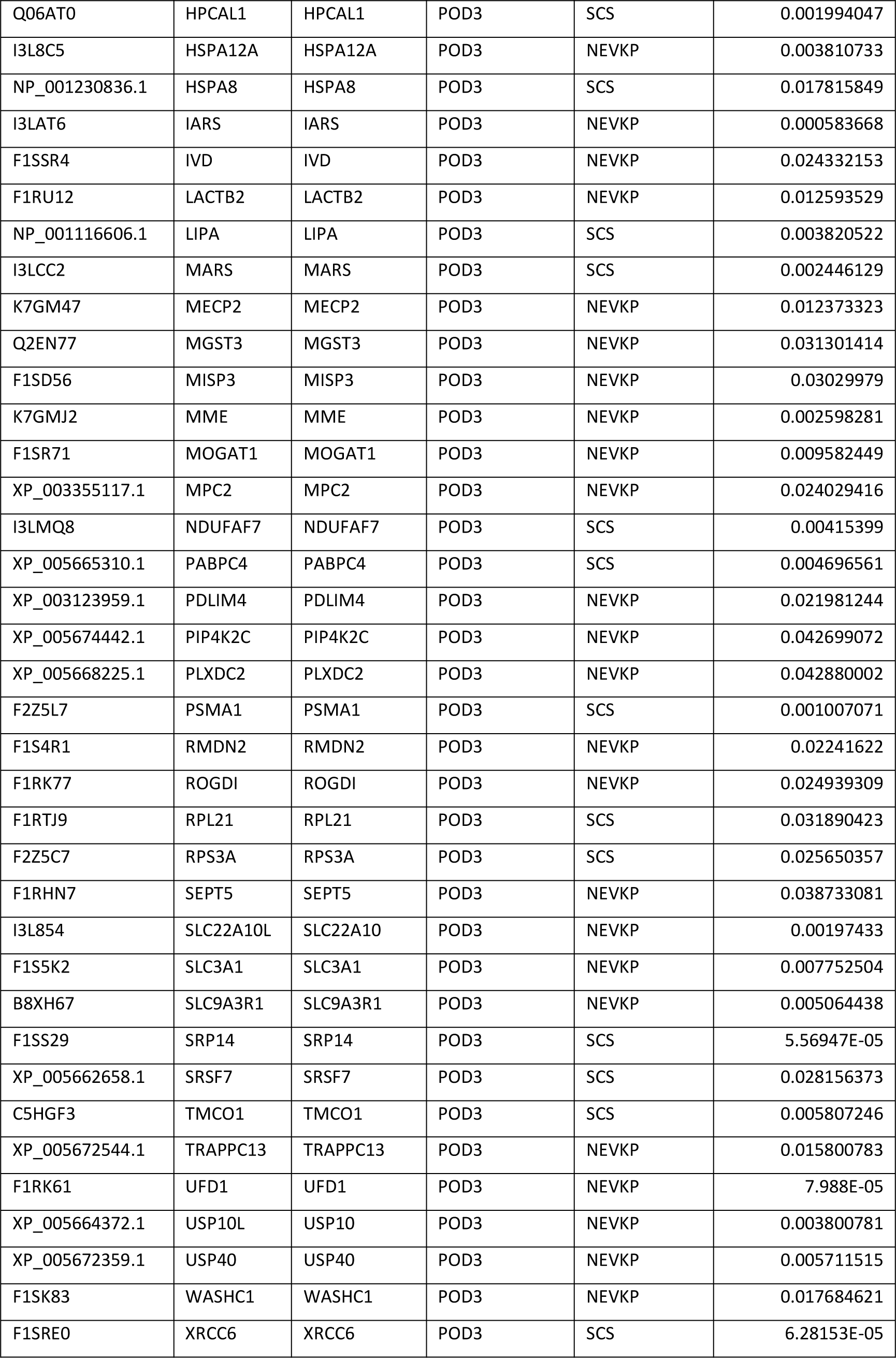
Details of the 70 proteins significantly differentially expressed between groups and across timepoints.

### Marked differences in the kidney proteome at POD3

To determine the changes in the kidney proteome over time following IRI associated with kidney transplantation, we performed an unsupervised clustering analysis using the binary tree-structured vector quantization (BTSVQ) algorithm. BTSVQ generates a binary tree dendrogram, iteratively partitioning the dataset into two subsets and utilizes self-organizing maps (SOMs) for data visualization.^38^ The first level of the dendrogram segregated POD3 samples from all others (Figure 2A). The SOMs of POD3 samples are markedly distinct from those at earlier timepoints, indicating that the expression profiles differ substantially between these groups of samples. Subsequent divisions of the dendrogram resulted in sub-clusters enriched for NEVKP-or SCS-samples respectively, but a further clear separation between the groups and/or timepoints was not evident. Supporting this observation, the majority (66/70) of DE proteins showed significant differences in expression between the experimental groups at POD3, while 4/70 DE proteins had significantly altered expression between groups at 30minutes post-reperfusion (Figure 2B, Table 1). The imputed (Figure 2B) and non-imputed (**Supplemental Figure 2**) heatmaps clustered the proteins similarly.

**Figure 2:**
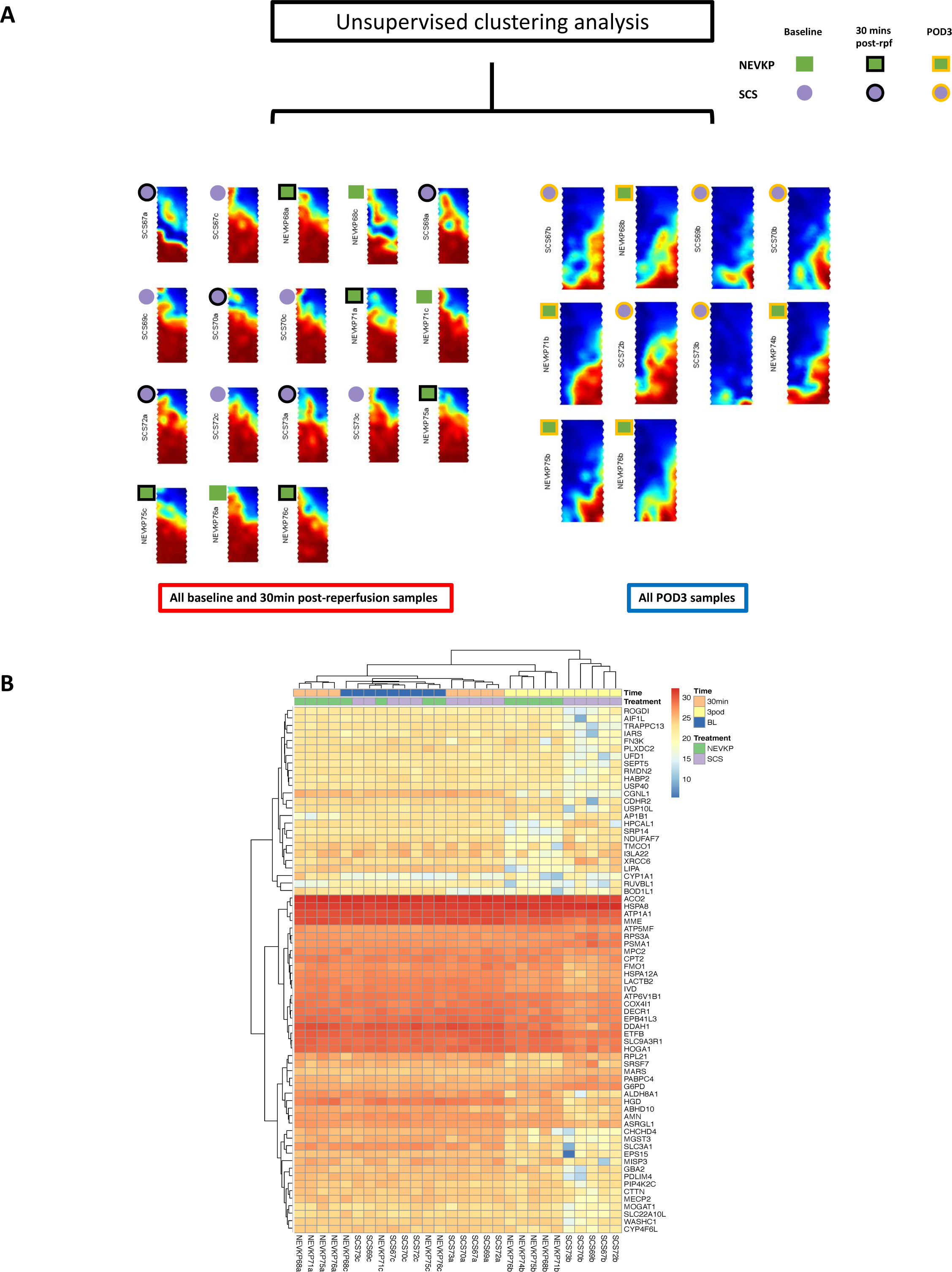
Expression profiles of the whole dataset, and of differentially expressed proteins show greatest differences between groups at POD3. (**A**) The first level of the binary-tree generated by BTSVQ clustering of the non-imputed protein expression profiles from each study sample, with SOM component planes at the nodes. Clear differences in the profiles of the POD3 samples versus the samples from the earlier timepoints are evident, accounting for the first binary division. NEVKP is represented by green squares, and SCS by purple circles. A black outer band denotes samples from 30 minutes post-reperfusion ("30 mins post-rpf"), and a gold outer band denotes POD3 samples. (**B**) Expression of the DE proteins across all samples depicted by heatmap with unsupervised hierarchical clustering of the proteins and samples. Columns represent each sample, and rows represent the differentially expressed proteins. The colour scale indicates the LFQ abundance of the protein across all samples ranging from blue (lower abundance) to red (higher abundance). Annotation of the columns details the experimental group and timepoint. BTSVQ, Binary tree-structured vector quantization; LFQ, normalized label-free quantification; POD3, post-operative day 3.

### GO and Pathway analysis

53/70 differentially expressed proteins were increased in NEVKP and 17 were increased in SCS (Table 1). We identified the significantly over-represented Gene Ontology terms among NEVKP-dominant and SCS-dominant proteins using g:Profiler.^39^ The most significant biological processes enriched in NEVKP-dominant proteins related to metabolism, specifically organic acid, amino acid and fatty acid/ lipid metabolism, and mitochondrial function (Figure 3A, **Supplemental Table 1**). Similarly, pathways significantly enriched among NEVKP-dominant proteins centred on metabolism, specifically, the TCA cycle and electron transport chain (Figure 3A), as determined by pathDIP.^40^ In contrast, SCS-increased proteins were annotated with biological processes relating to RNA catabolism and translation (Figure 3B, **Supplemental Table 2**).

**Figure 3:**
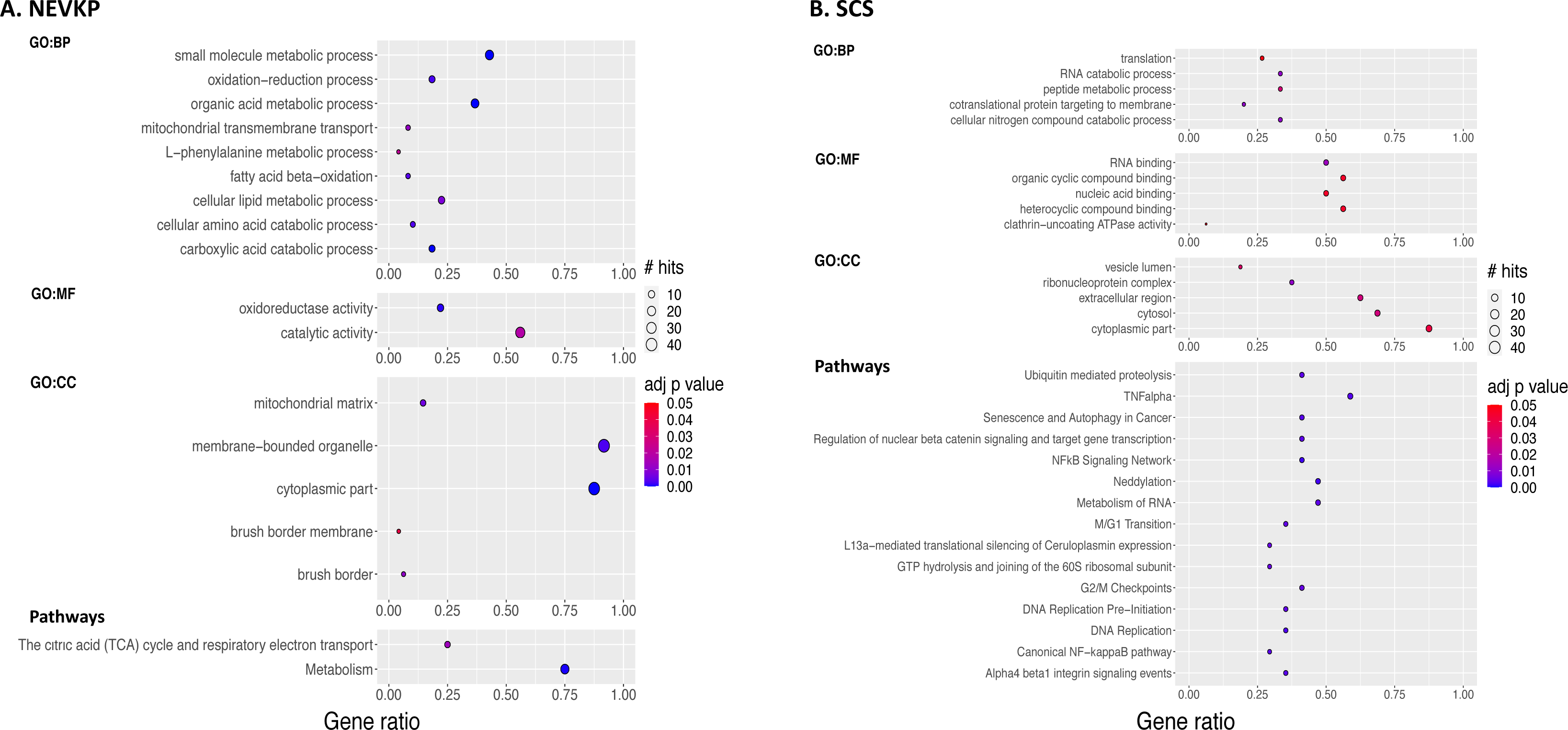
Gene ontology and pathway analysis of dysregulated proteins. The gene ontology terms significantly (BH-adjusted FDR <0.05) enriched among NEVKP-increased proteins (**A, left**) and SCS-increased proteins (**B, right**) respectively. The biological pathways (literature and experimentally proved protein-protein interactions) significantly enriched (BH adjusted FDR <0.05) among NEVKP-increased (**A,bottom left**) and SCS-increased (**B, bottom right**) proteins respectively. Node colour depicts the BH-adjusted FDR as shown by the colour bar; node size denotes the number of our DE proteins participating in the process/pathway in question, as shown by ‘# hits’; the x axis depicts ‘gene ratio’ which reflects a ratio of the number of DE proteins associated with that term: the number of DE proteins queried. BH-adjusted FDR, Benjamini-Hochberg adjusted false discovery rate; DE, differentially expressed; NEVKP, normothermic ex vivo kidney perfusion; SCS, static cold storage.

Consistent with the GO analysis, pathways related to DNA replication and RNA metabolism were significantly enriched among SCS-dominant proteins (Figure 3B, **Supplemental Table 3**). Furthermore, inflammation (TNF-α and NF-kB),^43^ integrin signalling (possibly mediating cell motility, and extracellular matrix organization^44^), and cell cycle arrest (reported following IRI,^45^ and linked with inflammation and fibrogenesis^46^) were significant among SCS-dominant proteins.

### Validation of findings using external datasets

We examined our findings in relation to other relevant datasets (Figure 4A, Table 2). We selected high-throughput studies relating to renal IRI^47–49^. Importantly, Damman et al.^49^ incorporates a cold ischemia component, analogous to SCS. As the kidneys and heart are metabolically similar^50^ we included a cardiac IRI^51^ study. We also included studies profiling other forms of kidney injury, specifically, septic-AKI,^52^ and CKD.^53^ We identified significant overlaps of our differentially expressed proteins with differentially expressed genes/proteins in the Port,^51^ Tran,^52^ Kang,^53^ Damman,^49^ and Huang^48^ datasets respectively (Figure 4A).

**Table 2:** External studies used for validation

| First Author | Year | Ref. No. | Organ | Organism | Specific context | Additional Details | Analysis of |
| --- | --- | --- | --- | --- | --- | --- | --- |
| Liu | 2017 | 47 | Kidney | Mouse | AKI, and AKI-CKD transition | Serial profiling over 12 month period following severe bilateral IRI | Gene expression (RNA-seq) |
| Huang | 2018 | 48 | Kidney | Rat | AKI-IRI | Analysis of affected and contralateral kidneys at 4 and 24 hours | Proteome |
| Damman | 2015 | 49 | Kidney | Human | Pre- and Post-Transplant | Peri-donation, post-cold ischemia, and post-reperfusion biopsies in LD, DCD and DBD donor kidneys | Gene expression (microarray) |
| Port | 2011 | 51 | Heart | Mouse | Myocardial Infarction | Biopsies from adjacent, non-infarcted left ventricle (or sham) at 2 days, 2 weeks and 2 months | Gene expression (microarray) |
| Tran | 2011 | 52 | Kidney | Mouse | AKI-Septic | LPS-induced AKI. Included profiles of groups with recovery and non-recovery of renal function | Gene expression (microarray) |
| Kang | 2015 | 53 | Kidney | Human | CKD | Microdissected tubulointerstitial samples, control v CKD (HTN or DM) | Gene expression (RNA-seq) |

**Figure 4:**
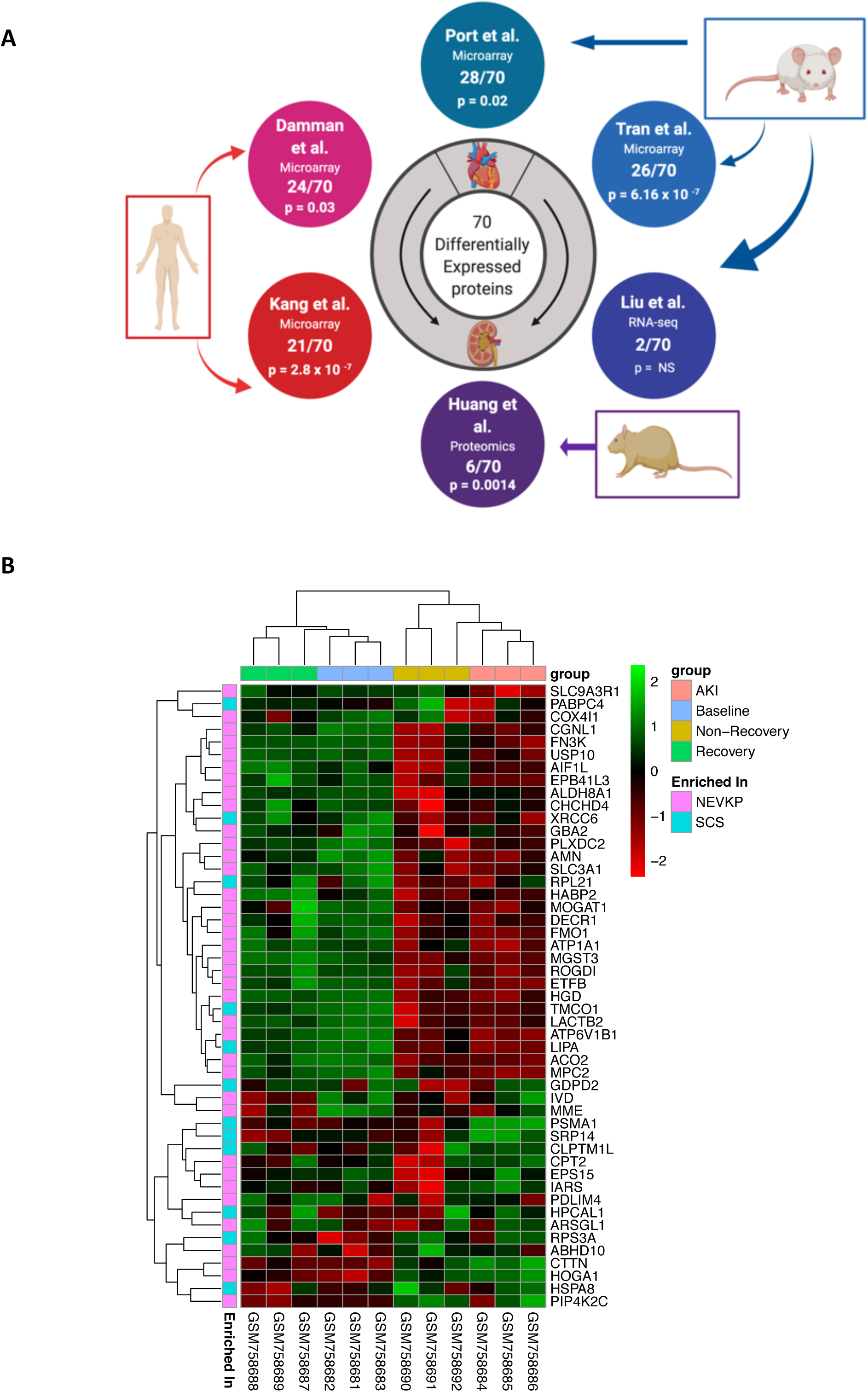
Validation of proteomics findings in external datasets. (**A**) We compared our list of DE proteins to the genes and proteins DE in a number of related studies derived from human (Damman et al., Kang et al.), mouse (Port et al., Tran et al., Liu et al.), and rat (Huang et al.) samples, as depicted. The overlap with specific DE proteins in our study for each external study is indicated. The significance of overlap was assessed using the hypergeometric test, with resultant p-values shown. (**B**) 49/70 of our DE proteins were represented in a mouse dataset of septic-AKI (Tran et al). The heatmap depicts the expression of these 49 proteins at the gene-level in the mouse dataset, using unsupervised hierarchical clustering. Columns represent the samples, and rows represent the genes, with relative expression of each gene across all samples demonstrated by pseudocolour scale ranging from −2 (red = lower expression) to +2 (green= higher expression). The columns are annotated to denote the experimental group of the mice in the Tran study. Annotation of the rows denotes increased expression in NEVKP or SCS respectively in the proteomic dataset. AKI, acute kidney injury; DE, differentially expressed; NEVKP, normothermic ex vivo kidney perfusion; SCS, static cold storage.

Predominantly, expression in NEVKP opposed the perturbation observed in disease or injury. **Supplemental Table 4** contains full lists of overlapping targets from each study. A subgroup of ^47^ differentially expressed proteins accounted for the overlap across studies (overlapping with ≥1 external study, the expression change in NEVKP opposing that observed in injury). The study by Tran et al^52^ permitted examination of our proteins in septic-AKI model that featured groups of mice with and without recovery of kidney function. 49/70 proteins had corresponding genes in the mouse microarray. We examined the expression of these 49 genes in the mouse dataset with unsupervised hierarchical clustering of genes and samples (Figure 4B). Significantly, these 49 proteins clearly separated those mice who recovered kidney function from those who did not. Mainly, the expression patterns of the proteins in NEVKP mirrored that observed in the mice at baseline and upon recovery of kidney function.

### Upstream Regulators

Our analysis suggested that preservation of key mitochondrial metabolic processes such as fatty acid oxidation (FAO) and TCA cycle /ATP-synthesis underpinned the proteome changes observed with NEVKP. The peroxisome proliferator-activated receptors (PPARs) and their transcriptional coactivator PPAR-γ coactivator-1α (PPARGC1A) are viewed as the key transcription factors regulating the expression of genes involved in fatty acid metabolism and mitochondrial biogenesis. Multiple sources of evidence implicate PPARs and PPARGC1a as potential upstream regulators in our dataset. A significant overlap exists (Figure 4A) between our differentially expressed proteins and the differentially expressed genes of datasets where PPARs and PPARGC1A were identified as key regulators (**Supplemental Tables 4-5**).^52–54^ Furthermore, using ARCHS4^41^ which integrates ChIP-seq data with large-scale RNA-seq data to predict transcription factor regulators of target genes, we verified that PPARG, PPARA, PPARD and/or the retinoid receptor X (RXR)-the common homodimer partner for ligand-bound PPAR signalling,^55,56^ were among the top-ranking transcription factors predicted to regulate 27/70 of our differentially expressed proteins (**Supplemental Tables 6-7**). Finally, using CATRIN, an extended transcription factor database which integrates the findings of multiple stand-alone transcription factor databases, we demonstrated that PPAR and RXR family members were predicted to regulate 65/70 differentially expressed proteins (Figure 5, **Supplemental Table 8**).

**Figure 5:**
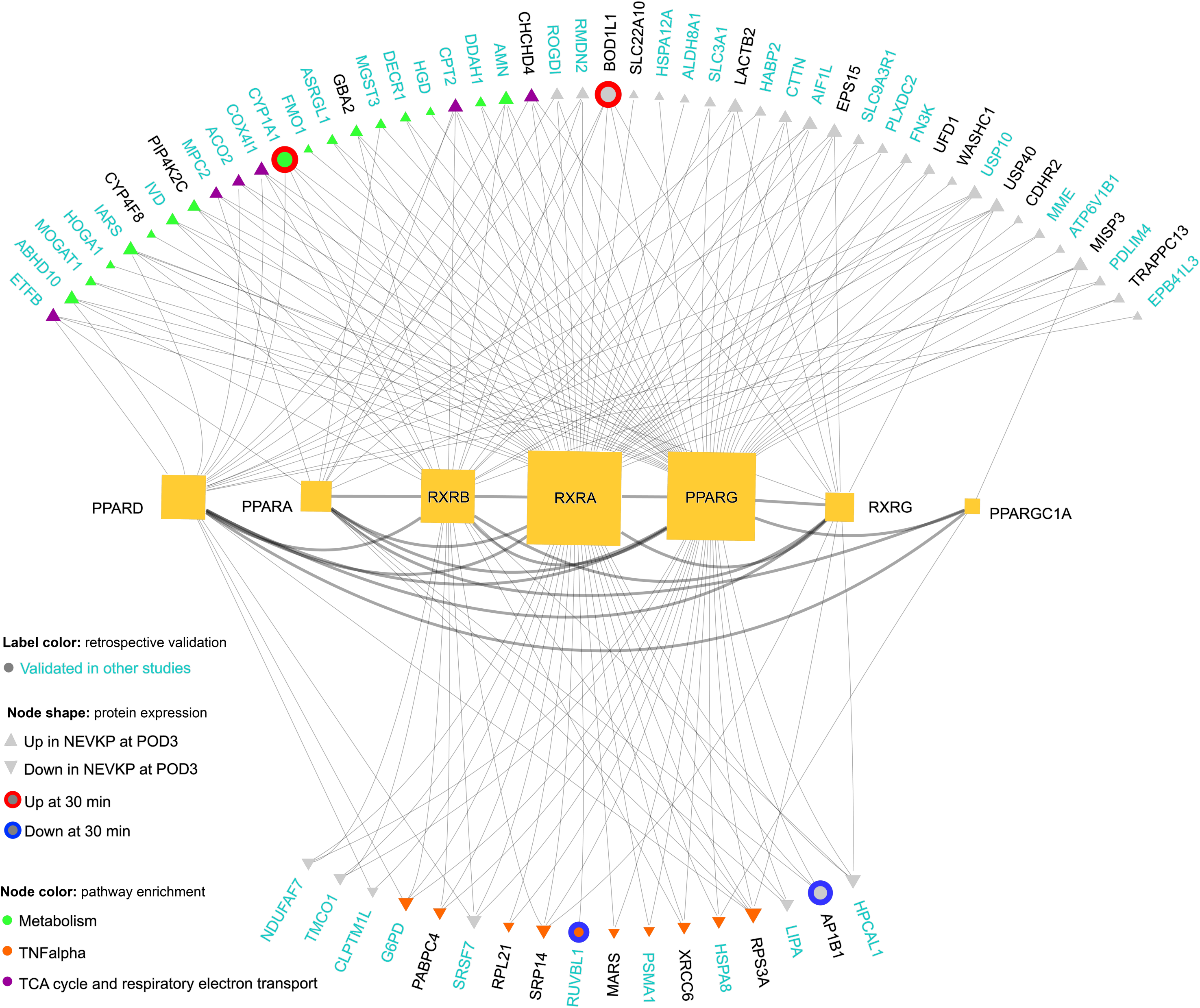
Regulatory interactions of PPAR family members and their coactivator (PPARGC1A) and signaling partners (RXR-family members) with our DE proteins. The regulatory interactions (‘grey lines’) of PPAR family transcription factors, PPARGC1A (co-activator) and RXRs with the DE proteins in our dataset were explored using an integrated transcription factor database, CATRIN. The network image was created using the NAVIGATOR software. The size of each transcription factor node corresponds to the number of our DE proteins regulated. Among the 70 DE proteins: ‘red’ and ‘blue’ outer circles denote increased and decreased expression in NEVKP at 30 minutes post-reperfusion respectively; ‘grey arrowheads’ reflect increased or decreased expression in NEVKP at the POD3 timepoint respectively. Nodes are then colored to indicate relevant pathway enrichments associated with the respective proteins. Finally, ‘cyan labelling’ indicates those proteins which were validated in independent datasets. DE, differentially expressed; NEVKP, normothermic ex vivo kidney perfusion.

### Experimental validation of key findings

Given the prominence of metabolic proteins in our dataset, we selected ETFB, CPT2 and COX4I1 for further validation. Consistent with the proteomics findings, ETFB and CPT2 were significantly increased in POD3 NEVKP-treated kidneys in comparison to SCS-treated kidneys (Figure 6A-B, Supplemental Figure 3). Immunohistochemical analysis of COX4I1 revealed more intense staining in the tubules of NEVKP-treated kidneys, compared to SCS (Figure 6C). Relative quantification of the stain confirmed this trend. We next validated our differentially expressed proteins at mRNA level. Among the proteins showing significant differences at 30minutes post-reperfusion, CYP1A1 had significantly increased gene expression in NEVKP mirroring the proteomics data (Figure 6D). Consistent with the proteomics data, COX4I1, MPC2, and ETFB showed significantly increased gene expression in NEVKP at POD3, while CPT2 expression demonstrated a similar trend (Figure 6E). There were no significant differences in expression of PPAR-family transcription factors at baseline between groups (**Supplemental Figure 4**), however, PPARA and PPARGC1A showed increased expression in NEVKP at 30minutes post-reperfusion (Figure 6F). PPARGC1A may mediate some of its reno-protective effects by augmenting expression of the lysosomal biogenesis regulator TFEB,^57^ which was significantly increased in NEVKP at 30minutes post-reperfusion (Figure 6F). Furthermore, PPARGC1A, PPARA, PPARD, RXRA and RXRB showed significantly increased expression in NEVKP compared to SCS at POD3 (Figure 6G).

**Figure 6:**
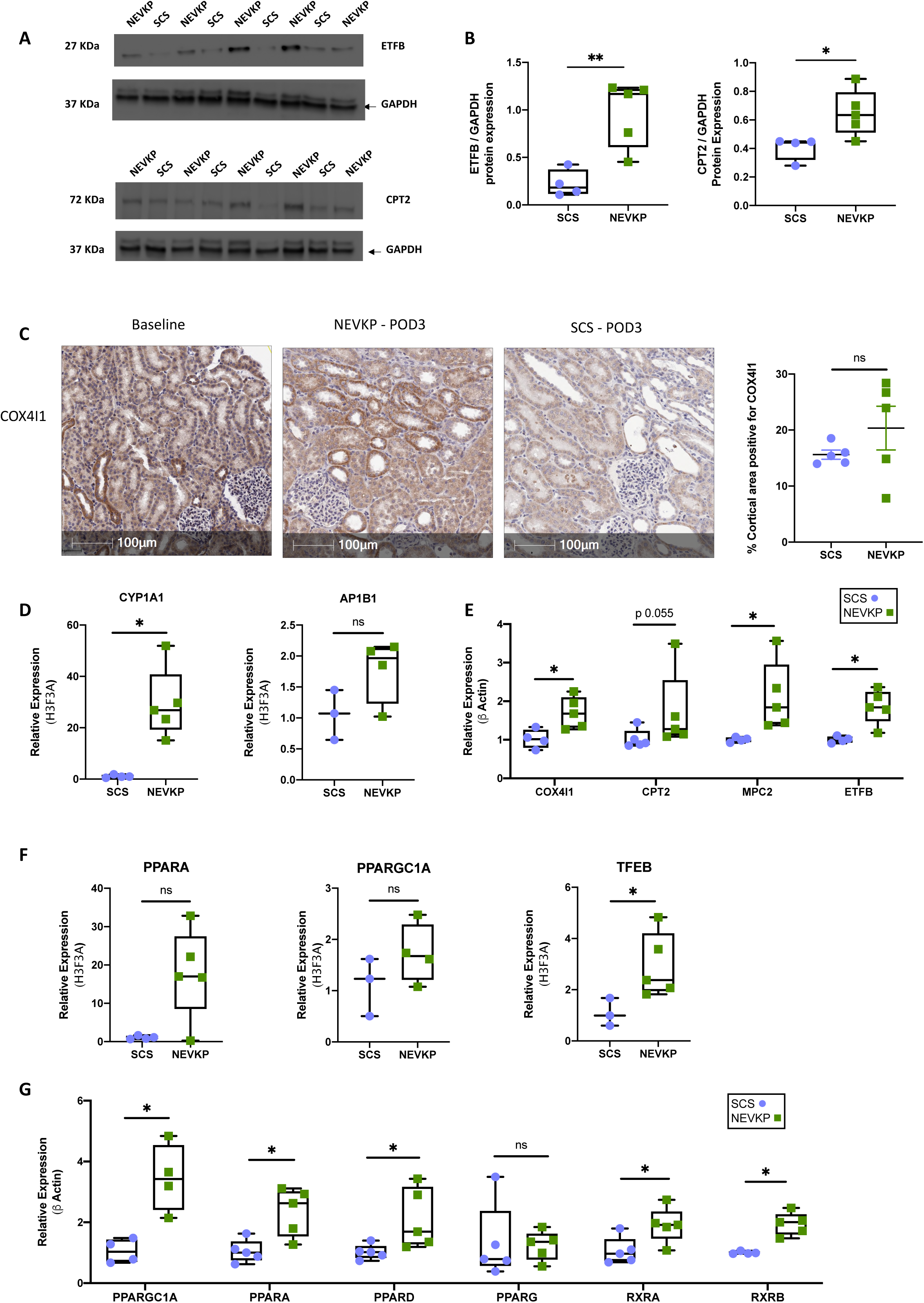
Validation studies of differentially expressed proteins and key findings. (**A-B**) Immunoblots representing ETFB, CPT2, and GAPDH protein expression in kidney biopsy tissue from the same animals used in the proteomics analysis. Intensities for ETFB and CPT2 were measured and normalized to GAPDH using Image J software. Mann Whitney test, n=–5 per group. (**C**) Expression of COX4I1 protein in NEVKP-and SCS-treated kidneys was verified by immunohistochemistry in new sections from POD3 formalin-fixed paraffin-embedded study samples. Magnification 20X. Scale bar 100μm. Mann Whitney test, n=5 per group. (**D**) Relative mRNA expression of AP1B1 and CYP1A1, in pig kidneys at 30minutes post-reperfusion (**E**) Relative mRNA expression of genes at POD3, related to the TCA cycle and DE in our dataset: COX4I1, MPC2, CPT2, and ETFB. (**F**) Relative mRNA expression of PPARA, PPARGC1A and TFEB in NEVKP and SCS groups at 30minutes post-reperfusion. (**G**) Relative mRNA expression of PPARGC1A, PPARA, PPARD, PPARG, RXRA and RXRB at POD3 in NEVKP and SCS groups. (**D-G**) Mann Whitney test, n=3–5 per group. *p<0.05, and **p<0.01 compared to SCS. DE, differentially expressed; POD3, post-operative day 3; NEVKP, normothermic ex vivo kidney perfusion; SCS, static cold storage; TCA, tricarboxylic acid.

### Urine metabolites

IRI engenders both early and sustained alterations in the metabolic profiles of kidney tissue, plasma, and urine^48,58,59^. We rationalized that NEVKP and SCS-induced changes in the proteome may be reflected in the urine metabolome. We conducted a targeted metabolomics analysis on urines collected from NEVKP and SCS at each timepoint. Firstly, we examined if the increased expression of CYP1A1 in NEVKP-treated kidneys or alternatively, the inflammatory pathways noted in the SCS-treated kidneys reflected activation of the aryl hydrocarbon receptor (AHR),^60^ as may occur secondary to accumulation of gut-derived uremic toxins (including indoxyl sulfate (IS), p-cresyl sulfate (pCS), p-cresyl glucuronide (pCG) and hippuric acid (HA)) in AKI^58,61^ and CKD.^62,63^ Secondly, given our hypothesis that PPARs and PPARGC1A represented likely upstream regulators of our NEVKP-related findings, we evaluated metabolites previously linked to the renoprotective effect of PPARGC1A (betaine, choline, carnitine and niacinamide) in IRI.^54^ Neither carnitine nor niacinamide were detected in our samples. For the analytes successfully measured, there were no significant differences in urinary excretion at baseline between groups (**Supplemental Table 9).** Urinary excretion of pCG and HA were significantly increased in SCS compared to NEVKP at POD3 (**Supplemental Figure 5a**). A similar (non-significant) trend was evident for IS (**Supplemental Figure 5a**). Urinary excretion of choline and betaine was increased in NEVKP compared to SCS at POD3, albeit not significantly (**Supplemental Figure 5b**).

Lactate and glucose are among the metabolites increased in the urine,^59^ and altered in tissue^58^ following IRI. At POD3, we observed increased urinary lactate and glucose in the SCS-treated group compared to NEVKP (**Supplemental Figure 5c-d**), as observed in prolonged DGF in a cohort of DCD-transplant recipients.^64^

## Discussion

This study was designed to uncover the molecular mechanisms underlying the beneficial effect of NEVKP. Our unique proteomics dataset profiles the molecular response to NEVKP and SCS following a DCD-type injury. There are three major findings: 1) proteins involved in mitochondrial energy production were significantly increased in NEVKP compared to SCS; 2) these proteins are significantly repressed in kidney disease of diverse etiologies as assessed in 6 external datasets; 3) PPAR and RXR transcription factors were computationally predicted upstream regulators of our metabolic proteins, and our gene expression findings support their increased activity in NEVKP.

We were struck by the observation that the differences between NEVKP-and SCS-proteomes at 30minutes post-reperfusion were minor, as shown by two independent analyses. This could be explained by insufficient time to cause changes in protein translation, most changes occurring in the low abundance proteome (typically undersampled), or that differences in response to the intervention are not driven by proteome changes at these early time points.

Our differentially expressed proteins featured critical enzymes governing mitochondrial energy metabolism. Proximal tubular epithelial cells (PTECs) utilize FAO as their preferred energy source, with inhibition of FAO associated with ATP depletion, intracellular lipid deposition, and cell death.^53^ PTEC lipid accumulation occurs in both AKI^54,65,66^ and CKD^53,67^ and results in reduced oxidative phosphorylation, generation of reactive oxygen species, and kidney fibrogenesis.^68^ Fatty acids must conjugate with carnitine to enter the mitochondria and consequently the carnitine phosphoryltransferase enzymes (CPT1 and CPT2) represent rate-limiting enzymes of FAO.^69^ Of the two, CPT2 is particularly vulnerable in IRI.^70^ ETFB is the β–subunit of the electron transfer flavoprotein which transfers electrons to the mitochondrial respiratory chain as FAO proceeds.^71,72^ Transcriptional repression of ETFB in ischemic cardiomyopathy is described.^73^ Suppression of mitochondrial transcripts in proportion to the degree of kidney dysfunction is also described in other AKI models.^52^ While FAO likely represents the primary means of ATP synthesis in PTECs, utilization of alternative substrates is described,^74,75^ with some evidence for a glycolytic shift following IRI.^76^ Moreover, other metabolically active segments of the kidney have alternative substrate preferences for ATP synthesis.^74,75^ Pyruvate, a hub metabolite for many metabolic pathways, enters the mitochondria via the mitochondrial pyruvate carrier (MPC), comprising two proteins (MPC1 and MPC2). Like PTECs, cardiomyocytes predominantly use FAO to generate ATP.^77^ Enhanced expression of MPC is seen in surviving myocardium post-ischemia, and may mediate tissue viability in this setting.^78^

The kidneys are highly metabolically active,^50^ requiring ATP for active solute transport against electrochemical gradients. Thus, normal kidney function is inextricably linked with mitochondrial energy production.^75,79,80^ These high energy demands may render the kidney especially vulnerable to ischemia.^54,81^ We propose that preserved expression of mitochondrial metabolic enzymes in NEVKP may underpin the improved kidney outcomes observed.

CYP1A1 was increased in NEVKP at 30minutes post-reperfusion-as reported after a similar *ex-vivo* perfusion period in lungs.^82^ The AHR is a prominent transcriptional regulator of CYP1A1, ^60^ and is potently activated by gut-derived protein-bound uremic toxins which accumulate in plasma and tissues in AKI and CKD.^62,63,83,84^ This activation is linked with the vascular dysfunction and systemic inflammation of CKD.^62,85–87^ In our study, these toxins were increased in urine of SCS pigs, potentially linking to the inflammatory pathways of SCS. AHR-independent pathways also regulate CYP1A1 expression^60,88–90^ including PPARA.^91^ CYP1A1 has well-described roles in drug metabolism and lipid oxidation.^89^ CYP1-enzymes participate in the oxidative biosynthesis of polyunsaturated fatty acids,^92^ and the specialized pro-resolving lipid mediators (SPMs) derived from these precursors.^93^ SPMs actively coordinate the resolution of acute inflammation thereby limiting the inflammatory response.^94,95^ Analysis of peritonitis-associated lipid-mediator metabolomes in CYP1-family knockout mice revealed increased neutrophil recruitment, elevated leukotrieneB4, and reduced intermediary compounds of SPM biosynthesis.^96^ The induction of CYP1A1 in NEVKP may reflect these non-classical, pro-resolving pathways of activation.

PPAR-family members and their transcriptional co-activator PPARGC1A emerged as likely upstream regulators in our dataset, with PPARA showing increased expression at 30minutes post-reperfusion in NEVKP, and PPARA/D, RXRA/B and PPARGC1A showing significantly increased expression in NEVKP at POD3. The renoprotective effects of PPARGC1A, particularly, have been described in models of septic,^52,97^ toxic,^57,98^ and ischemic^54,99,100^ AKI. Downregulation of PPARGC1A and related transcripts is observed in CKD of diverse etiologies,^53,101^ and implicated in the development of inflammation^102^ and age-related fibrosis in the kidney.^103^ Kidney transplants with increased PPARGC1A expression demonstrated a faster and more complete recovery from DGF.^104^ PPARGC1A is considered the ‘master regulator’ of mitochondrial biogenesis, binding to a host of transcription factors (most notably PPAR-family members) to increase expression of genes that augment mitochondrial abundance, oxidative phosphorylation, and FAO.^105–108^ Observations that tubular PPARGC1A can reduce the severity of AKI and accelerate functional resolution^54,57,99^,^109^ are consistent with the high metabolic activity of PTECs.^110^ Less metabolically active kidney cell types including endothelial cells54 and podocytes^111,112^ may not experience the same benefit, suggesting a cell-type specific role for PPARGC1A in the kidney.

Previous observations about the metabolic footprint of PPARGC1A renoprotection^54^ prompted us to examine related markers in the urine. A modest increase in urinary choline was evident in the NEVKP-treated group. Choline and betaine are renal osmolytes113. Increased urinary osmolytes are reported following cold ischemia, and hypothesized to reflect medullary cell damage.^114^ Increased urinary betaine and choline are reported in CKD,^115^ and incipient diabetes.^116^ Conversely, other evidence suggests our observed increases in urinary choline could reflect increased PPAR activity.^54^ Increased concentrations of choline are noted in kidneys of wild-type mice in comparison to PPARA-/-mice.^117^ Treatment of healthy individuals with fibrates (PPARA-agonists) results in increased urinary choline and betaine,^118^ with similar findings in animal models.^119^ Our urinary observations support our proteomic and gene level findings which together suggest that the alterations observed in NEVKP-treated kidneys may reflect increased PPARA and PPARGC1A activity.

Our study has many strengths. Given the anatomical and physiological similarities of pigs and humans, our large animal model is readily clinically translatable, and well-suited to the study of IRI and transplantation. In contrast to previous studies,^82,120^ we assess the impact of NEVKP post-transplant, and examine the functional significance of *ex-vivo* observations. Our systems biology approach incorporates transcriptomic and targeted metabolomic analyses, as well as an analysis of upstream regulators. Finally, this is a novel dataset; to our knowledge, this is the first proteomics study related to NEVKP. Notwithstanding the strengths of our study, some limitations exist. Our porcine DCD-model lacks some elements typically observed clinically, most notably, severe antecedent illness in the donor, alloantigen exposure, and post-operative immunosuppression. The structural and functional annotation of the pig genome remains incomplete,^121^ rendering biological interpretation challenging. Lastly, while our proteins were predicted to be regulated by PPAR/RXR transcription factors, which was supported by their alteration at mRNA level, it is plausible that post-translational modifications contributed to differences in protein abundance.

In summary, we present a detailed analysis of the changes in the kidney proteome induced by NEVKP in comparison to SCS. We conclude that preservation of key mitochondrial enzymes mediating crucial metabolic pathways may be responsible for the superior kidney outcomes seen with NEVKP and that these effects are coordinated by the PPAR-family of transcription factors and their co-activator PPARGC1A (Figure 7). Our findings suggest potential therapeutic targets to ameliorate IRI in kidney transplantation.

**Figure 7:**
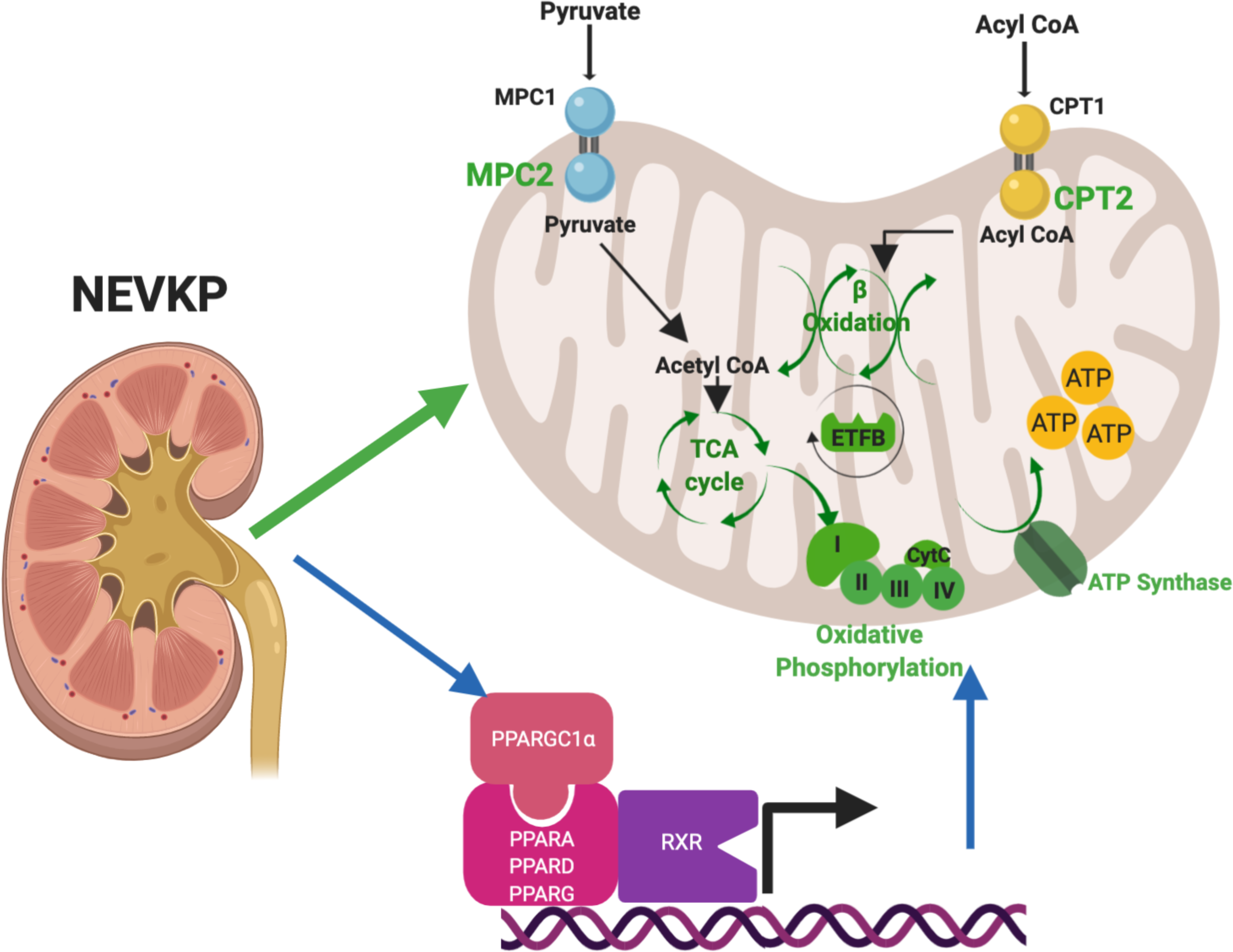
Proposed role of NEVKP in attenuating ischemia-reperfusion injury in a DCD-model of auto-transplantation. NEVKP is associated with preserved expression of proteins mediating critical metabolic processes in the mitochondria in comparison to SCS. We demonstrate increased expression of proteins mediating the entry of key energy-producing substrates into the mitochondria (MPC2, CPT2), proteins involved in the TCA cycle (ACO2), electron transfer (ETFB), oxidative phosphorylation (COX4I1), and ATP synthesis (ATP5MF) resulting in enrichment of fatty acid ≥-oxidation, the TCA cycle and oxidative phosphorylation. All NEVKP-increased processes are shown in green. The blue arrows represent our findings on gene expression that these effects are centrally regulated by members of the PPAR-family of transcription factors (PPARA, PPARD, PPARG), RXRA, RXRB and their transcriptional co-activator PPARGC1a. DE, differentially expressed; NEVKP, normothermic ex vivo kidney perfusion; SCS, static cold storage; TCA, tricarboxylic acid.

## Author Contributions

AK conceived the study. AK, MS, and LAR participated in study design; CMcE, SC-F, SR, IB, JMK, PU, AAERP, SF, JV, BLU, MS and AK carried out experiments; CMcE, SC-F, TT, CP, RJ, IJ and AK analyzed the data; CMcE, CP, RJ, and TT made the figures; CMcE, SC-F and AK drafted and revised the paper; all authors approved the final version of the manuscript.

## Acknowledgements

AK is supported by a Kidney Foundation of Canada operating grant, the Kidney Research Scientist Core Education and National Training (KRESCENT) program, Kidney Foundation of Canada Predictive Biomarker Grant, CIHR Catalyst Grant, and Canada Foundation for Innovation. She has also received funding from the Toronto General Hospital Research Foundation and the Multi-Organ Transplant program. CMcE is supported by the Menkes fellowship, and a University Health Network Multi-Organ Transplant fellowship. SC-F is supported by the KRESCENT program. IJ, TT & CP were supported in part by Ontario Research Fund (#34876), Natural Sciences Research Council (NSERC #203475), Canada Foundation for Innovation (#29272, #225404, #30865), Krembil Foundation and IBM. BLU is supported by a Canada Foundation Innovation award. LAR is supported by the Hospital for Sick Children Transplant and Regenerative Medicine Centre.

## Data Availability

The data supporting the findings of this study are have been deposited to the ProteomeXchange Consortium (http://proteomecentral.proteomexchange.org) with the dataset identifier PXD015277.

## Supplemental Material Table of Contents

### Supplemental Methods

#### Supplemental Figure Legends

##### Supplemental Figures

SF1: Distribution of protein abundance with QRILC-imputed values

SF2: Heatmap depicting the expression profiles of the DE proteins using non-imputed data

SF3: Uncropped western blot images

SF4: Gene expression of PPAR-family and related transcription factors at baseline

SF5: Analysis of Urine Metabolites

##### Supplemental Tables

ST1: GO analysis of NEVKP-dominant proteins

ST2: GO analysis of SCS-dominant proteins

ST3: Pathway analysis (pathDIP)

ST4: Summary table of the validation with external datasets

ST5: Overlap of DE proteins with PPARGC1A-regulated IRI dataset

ST6: ARCHS4 analysis-Ranking of top transcription factors

ST7: ARCHS4 analysis-z scores of PPAR-family transcription factors in genes encoding DE Proteins

ST8: Network mapping using CATRIN relating to Figure 5.

ST9: Urinary metabolite concentrations adjusted by urinary creatinine ST10: Primer sequences used in RT-PCR

ST10: Primer sequences used in RT-PCR

