## Supplemental Methods and Figures for "Normothermic Ex-vivo Kidney Perfusion in a Porcine Auto-Transplantation Model Preserves the Expression of Key Mitochondrial Proteins: An Unbiased Proteomics Analysis"

### Supplemental Material Table of Contents

**Supplemental Methods**

**Supplemental Figure Legends**

**Supplemental Figures**

SF1: Distribution of protein abundance with QRILC-imputed values

SF2: Heatmap depicting the expression profiles of the DE proteins using non-imputed data

SF3: Uncropped western blot images

SF4: Gene expression of PPAR-family and related transcription factors at baseline

SF5: Analysis of Urine Metabolites

**Supplemental Tables**

ST1: GO analysis of NEVKP-dominant proteins

ST2: GO analysis of SCS-dominant proteins

ST3: Pathway analysis (pathDIP)

ST4: Summary table of the validation with external datasets

ST5: Overlap of DE proteins with PPARGC1A-regulated IRI dataset

ST6: ARCHS4 analysis- Ranking of top transcription factors

ST7: ARCHS4 analysis- z scores of PPAR-family transcription factors in genes encoding DE Proteins

ST8: Network mapping using CATRIN relating to Figure 5.

ST9: Urinary metabolite concentrations adjusted by urinary creatinine

ST10: Primer sequences used in RT-PCR

### Supplemental Methods

##### Experimental model

As previously described,[1,2] 3 month old male Yorkshire pigs were used in this model. Following induction of general anesthesia, the right renal artery and vein were clamped for 30 minutes, mimicking a DCD-type injury. Following this, the right kidney was removed, and the vessels were cannulated and flushed with 400-mL histidine-trytophan-ketoglutarate. The right kidney was subjected to either 8 hours of SCS or 8 hours of continuous pressure-controlled NEVKP, followed by auto-transplantation. Prior to re-implantation, the contralateral kidney was removed. The pigs were followed up for 3 days following transplantation, with daily assessment of renal function, before being euthanized. The study was approved by the Animal Care Committee of the Toronto General Research Institute, Ontario, Canada. All animals received humane care in compliance with the "Principles of Laboratory Animal Care" formulated by the National Society for Medical Research.

##### NEVKP

Normothermic ex vivo perfusion was conducted as previously described.[1-3] The perfusion circuit is based on modified neonatal cardiopulmonary bypass technology. A centrifugal pump propels the perfusion solution into the oxygenator, where it is oxygenated and warmed to 37 °C. After passing the arterial filter (95% $O_2$, 5% $CO_2$, 2L/min), the perfusate is driven with a pressure of 65 mm Hg through the renal artery into the graft which is housed in a chamber. The venous outflow leads the perfusate back into the venous reservoir. Any urine produced is collected throughout the perfusion. As previously described, the perfusion circuit was primed with Ringer's lactate (200mL), Steen solution TM (150mL, XVIVO Perfusion AB, Goteborg, Sweden), washed leukocyte-filtered erythrocytes (125mL), double reverse osmosis water, 8mL sodium bicarbonate (8.4%), 1.8mL calcium gluconate (10%, 100 mg/mL), and heparin to achieve a near physiologic perfusate composition.[4] During perfusion, urine output was replaced with Ringer's lactate. Verapamil (0.25 mg/h) was administered intra-arterially, and amino acids and glucose (1 mL/h) with insulin (5 IU/H) were administered continuously during the perfusion and were adapted to maintain perfusate glucose level between 5 and 15 mmol/L. Blood gas parameters were measured hourly, with optional administration of bicarbonate to maintain acid-base homeostasis, though this was typically not required after the first hour of perfusion.[3]

**Sample collection and storage**

18G core biopsies were taken at the first two timepoints. At POD3 a wedge biopsy was reserved. Urine samples obtained from bladder puncture were immediately centrifuged (9,000*g* x 10 mins) and the supernatant reserved. All samples were snap-frozen in liquid nitrogen, then stored at -80°C.

**Sample preparation for proteomics analysis**

Frozen porcine kidney biopsy samples were covered with 0.1% Rapigest, followed by homogenization at 15,000rpm for 15-30 seconds on the Polytron PT3100 homogenizer. Samples were subsequently sonicated for 10 seconds, three times, on ice. They were then centrifuged at 15,000g at 4°C for 20 minutes. The supernatant was collected and vortexed. Total protein concentration was measured using Coomassie assay and each sample was normalized to 250μg of total protein. Two samples had significantly less than 100μg of total protein and were thus eliminated from further analyses. The remaining 28 samples underwent denaturation at 80°C for 15 minutes, reduction with 10mM DTT for 15 minutes at 65°C and finally, alkylation with 20mM iodoacetamide in the dark, at room temperature, for 40 minutes. The samples were then incubated overnight with trypsin (Promega) 1:50 w/w at 37°C. The following morning, TFA was added to each sample at 1% v/v. Each sample was vortexed for 1 minute, then left at room temperature for 5 minutes. The samples were subsequently centrifuged at 15,000*g* for 10 minutes. Supernatants were then transferred into new tubes and the pellets were discarded. Individual samples were resuspended in SCX mobile phase A (0.26 M formic acid in 5% v/v acetonitrile; pH 2-3) and loaded directly onto a 500 µL loop connected to a PolySULFOETHYL A™ column (2.1 mm ID × 200 mm, 5 µm,

200 Å, The Nest Group Inc., MA). SCX chromatography and fractionation were performed on an HPLC system (Agilent 1100) using a 60-min two-step gradient. An elution buffer which contained mobile phase A with the addition of 1 M ammonium formate was introduced at 10 min and increased to 20% at 30 min and then to 100% at 45 min. Fractions were collected every 1 min from the 20 min time point onwards. The resulting fractions corresponding to chromatographic peaks of eluting peptides were pooled into 7 fractions.

**Tandem mass spectrometry**

Peptides were identified by LC-MS/MS as described previously.[5] Peptides from each fraction were extracted with 10 µL OMIX C18 MB tips (Agilent, USA eluted in 3µL of 65% v/v acetonitrile, diluted to 40 µL with 0.1% v/v formic acid in pure water, and loaded onto a 3.3 cm C18 pre-column (with an inner diameter of 150 µm; New Objective), packed in-house with 5 µm Pursuit C18 (Agilent, USA). Eluted peptides from the trap column were subsequently loaded onto a resolving analytical PicoTip Emitter column, 15 cm in length (with an inner diameter of 75 µm and 8 µm tip, New Objective) and packed in-house with 3 µm Pursuit C18 (Agilent, USA). The columns were operated on the EASY-nLC system (Thermo Fisher Scientific, San Jose, CA), and this liquid chromatography setup was coupled online to Q-Exactive Plus mass spectrometer (Thermo Fisher Scientific, San Jose, CA) using a nano-ESI source (Thermo Fisher Scientific). Each fraction was run using a 60-min gradient and analyzed in data-dependent mode in which a full MS1 scan acquisition from 400-1500m/z in the Orbitrap mass analyzer (resolution 70,000) was followed by MS2 scan acquisition of the top 12 parent ions. The gradient was increased from 1% to 5% Buffer B at 2 minutes, followed by an increase to 35% Buffer B at 49 minutes, 65% at 52 minutes and 100% at 53 minutes. The following parameters were enabled: monoisotopic precursor selection, charge state screening and dynamic exclusion (45.0 seconds). In addition, charge states of +1, 5 – 8, >8 and unassigned charge states were not subjected to MS2 fragmentation. For protein identification and data analysis, XCalibur software v3.0.63 (Thermo Fisher) was utilized to generate RAW files of each MS run.

**Protein identification and quantification**

The raw mass spectra from each fraction were analysed using Andromeda search engine (MaxQuant software v.1.5.3.28) against the nonredundant Sus scrofa database generated from a nonredundant union of 26139 porcine sequences from UniProtKB, 24556 sequences from NCBI RefSeq databases and cRAP database of common contaminants (as previously published).[6] Reverse decoy mode was used. Tryptic peptides were selected with up to two mis-cleavages. Methionine oxidation and N-terminal protein acetylation were selected as variable modifications. Carbamidomethylation was selected as fixed modification. Protein and site FDR were set at 0.01. The top 12 peaks were selected for MS/MS. MS/MS parent tolerance was set to 20ppm and fragment tolerance was set to 0.5Da. The minimum ratio count was set to 1. Matching between runs was selected, with a matching time window of 0.7 minutes and an alignment window of 20 minutes. Label free quantification was performed and normalized protein LFQ intensities were used for subsequent analyses. The data were analysed using Perseus v.1.5.2.6. Reverse hits and contaminants were removed. Normalized LFQ intensities were log2-transformed and the samples were annotated according to the group (i.e. NEVKP or SCS) and time point (i.e. BL, 30-minutes post-reperfusion, POD3). We then filtered data to include only those proteins that were identified in at least 5 samples at any time point.
The mass spectrometry data have been deposited to the ProteomeXchange Consortium (http://proteomecentral.proteomexchange.org) via the PRIDE partner repository[7] with the dataset identifier PXD015277.

**Proteomic data analysis**

Missing values were imputed using the widely used QRILC method, which performs the imputation of left-censored missing data using random draws from a truncated distribution with parameters estimated using quantile regression with the R package imputeLCMD (v2.0) under default parametrization.[8,9] 300 independent replicates of the imputed data were created. For each, we performed 2-way ANOVA followed by Tukey's HSD test. The resultant p-values were adjusted for multiple testing by the FDR method. Finally, to obtain robust estimates of statistical significance, we calculated geometric mean of the adjusted p-values across the imputation replicates. Proteins whose p-value <0.05 for association with the effect of treatment, time, and their interaction term were depicted by heatmap with hierarchical clustering of proteins and samples.

For initial clustering and visualization of the dataset, we used Binary Tree-Structured Vector Quantization (BTSVQ) algorithm which iteratively generates a binary tree by partitioning the samples into two subsets at each level of a tree. The algorithm uses self-organizing maps (SOMs) to cluster biochemical variables, and for sample and cluster visualisation.[10] BTSVQ is available from (http://www.cs.toronto.edu/~juris/btsvq/overview.html), and is implemented using the SOM toolbox in Matlab (http://www.cis.hut.fi/projects/somtoolbox/).

Gene Ontology and pathway enrichment were calculated using g:Profiler[11] and pathDIP[12] respectively. The human orthologues of the genes encoding for the 70 differentially expressed proteins were used as an input for the Gene Ontology and pathway enrichment analysis. Default settings on g:Profiler [11] (https://biit.cs.ut.ee/gprofiler/gost) were used apart from the selection of Benjamini-Hochberg FDR 0.05 as the significance threshold, and the exclusion of electronic GO annotations. During pathway enrichment analysis using Pathdip[12] (v3) (http://ophid.utoronto.ca/pathDIP/), we selected the extended pathway associations, integrating core pathways with experimentally proven protein-protein interactions, with the default minimum confidence level for predicted associations of 0.99 accepted.

**Upstream regulator analysis and network visualization**

The ARCHS4 pipeline has aligned and curated the majority of published RNA-seq data (GEO/SRA) from human and mouse. The ARCHS4 web interface makes this information available at the gene and transcript levels. Additionally, the web resource enables exploration of the processed data through a number of querying tools, including the prediction of upstream transcription factors based on prior knowledge and co-expression with identified targets as determined by ChIP-seq data from the ChEA and ENCODE gene set libraries.  We interrogated ARCHS4[13] to identify transcription factors predicted with a high likelihood (i.e. the top 10 ranking factors with z-score >2) to regulate genes encoding our differentially expressed (DE) proteins. The resultant list of transcription factors was then ranked on the number of our DE proteins they were predicted to regulate (shown in **Supplemental Table 6 and 7**)

Catrin (http://142.150.188.233:9080/Catrin/index.jsp), is a **Ca**talogue of **T**ranscriptional **R**egulatory **In**teractions that integrates 15 separate transcription factor databases. Many of these databases utilize different methods to identify transcription factor to gene pairs, and some of the databases focus only on a single transcription factor or on a single class of transcription factors. For this reason, the overlap across databases is quite poor, and the integration of such data extends the coverage of transcriptional regulatory interactions. DE proteins were used to query Catrin with all resources except TF2DNA.experimental. Regulatory interactions were obtained from specified transcription factors (PPARA, PPARD, PPARG, PPARGC1A, RXRA, RXRB, RXRG) to query genes. Due to the completeness of Catrin data, interactions retrieved from it were used to build a network.  Network visualization analysis was performed using NAViGaTOR 3.0.10, a stand-alone scalable software designed to create, annotate, visualize and analyze networks.[14]

**Analysis of external datasets**

We explored the context of our findings in relevant external datasets by examining for overlap between our 70 DE proteins and the DE proteins or transcripts in the external studies (**Supplemental Table 4**).

Damman et al.[15] examined gene expression in deceased donor biopsies, at three timepoints: retrieval, after cold ischemia, and 45-60 minutes post-reperfusion. Living donor kidney biopsies at two timepoints: before clamping of the renal artery, and 45-60 mins post-reperfusion were also examined. Using the detailed annotation supplied with this study on GEO2R (GSE43974) we identified the genes DE (FDR-adjusted $p < 0.05$) at the post-reperfusion timepoint between DCD donors who experienced DGF, and living donors who did not (viewing this as the most analogous comparison to SCS- and NEVKP-kidneys respectively).

Tran et al.[16] studied the gene expression profiles of mice subjected to a septic-AKI induced by lipopolysaccharide. In this experiment, all mice received fluid resuscitation after developing the AKI; some then recovered baseline renal function, while others did not. Using the annotation details provided (GSE30576 - GPL8759) on GEO, we identified the genes DE (FDR adjusted $p < 0.05$) between mice with an AKI in comparison to baseline. Separately, we downloaded the raw data files for this experiment (GSE30576 -GPL8759), and after pre-processing and $Log_2$ normalization (Limma v3.34.14)[17], generated an expression dataset for all animals (Baseline, AKI, Recovered, and Non-recovered). We extracted the values for the genes corresponding to the 70 DE proteins from our proteomics dataset. In total, 49 of the proteins were represented in the murine dataset. The expression profiles of these genes across all samples was plotted using a heatmap (pheatmap v1.0.12) with unsupervised hierarchical clustering of genes and samples.

Kang et al.[18] performed RNA sequencing in micro-dissected tubulointerstitial samples to identify genes DE between individuals with CKD and healthy controls. Details of DE transcripts are supplied in their supplement.

Liu et al.[19] used RNA-seq at multiple timepoints to report the temporal-specific alterations in gene expression following bilateral severe IRI, and the ensuing progression to CKD. The list of genes DE in these mice when compared to controls, is supplied in their supplement. Huang et al[20] performed a LC-MS/MS proteomic analysis of rat kidney cortices at 4 and 24 hours post-reperfusion following a 45-minute period of unilateral IRI, using contralateral kidneys and healthy controls as comparators. The list of DE proteins is provided in their supplement.

Port et al[21] examined gene expression in adjacent but non-infarcted left ventricle of mice in a model of myocardial infarction at three timepoints (2 hours, 2 days and 2 weeks post-infarction), using sham-operated time-matched controls as controls. The list of DE genes identified in this study is provided in their supplement.

For each study, the statistical significance of the overlap identified was assessed using the hypergeometric test in R.

**Gene expression**

Total RNA was extracted from frozen wedge and core biopsies of pig kidney tissue using the RNAeasy Mini Kit (Qiagen). RNA concentration was measured in each sample using Nanodrop (Thermo Scientific), and 500ng of RNA were subsequently retrotranscribed to cDNA using the High Capacity cDNA Reverse Transcription Kit (Applied Biosystems). Gene expression of ACADM, ACADVL, AP1B1, ATP5PO, COX4I1, COX5B, CPT2, CYP1A1, ETFB, MPC2, PABPC4, PPARA, PPARD, PPARG, PPARGC1A, RXRA, RXRB, and TFEB was determined by real-time quantitative PCR using Power SYBR® Green PCR Master Mix (Applied Biosystems) in a StepOne Plus System (Applied Biosystems). For each time point, gene expression data were normalized to the most stable housekeeping gene across conditions: RPS16 (baseline), H3F3A (30 min), and ACTB (POD3). Primer sequences are summarized in **Supplemental Table 10**.

**Immunoblotting**

Fresh protein lysates were made from stored residual biopsy tissue. After protein quantification (Micro BCA Protein Assay kit (ThermoFisher Scientific)), 10µg of protein per well was loaded onto 10% acrylamide gels (BioRad) and separated by SDS-PAGE. Membranes were incubated with antibodies to CPT2 (26555-1-AP, Proteintech®), ETFB (LS-C81860 / 146214. GAPDH (CB-1001, Millipore Sigma) was used as a loading control.  Secondary antibodies included: HRP-conjugated anti-mouse (P044701-2, Agilent) and HRP-conjugated anti-rabbit (A0545, Sigma). Western blot images were acquired using the DNR Bioimaging Systems MicroChemi 4.2. which incorporates the molecular weight reference image with the image of the membrane.

Following detection, bands were quantified by densitometry using Fiji/Image J (1.x).[22,23] For validation, 4-5 animals per group were studied.

**Immunostaining**

For COX4I1 immunohistochemistry, antigen retrieval was performed in dewaxed and rehydrated 5μm sections by heating the samples with a citrate buffer (pH6) (Abcam) in a pressure cooker for 20 minutes. Sections were then incubated with rabbit anti-COX4I1 antibody (dilution 1:100, MA-15078, Invitrogen) diluted in Dako diluent (Dako, No.S0809). HRP- conjugated anti-rabbit (MP-7401, Vector Labs) was used as secondary antibody. Samples were further incubated with DAB, then counterstained with hematoxylin to visualize nuclei. Slides were digitally scanned in a ZEISS Axio Scan.Z1 system.

Cortical areas on the whole slide image (wedge biopsy, POD3) were assessed for positive staining. Areas of capsule, medulla, artefact and large blood vessels were excluded from analysis. To assess protein expression, the percentage of positive staining in the cortical area assessed was quantified using the Halo software (Indica Labs, version 3.0.311).

**Urine Biochemistry and Creatinine measurements:**

The BioProfile FLEX analyzer (novaBiomedical) was used to measure glucose and lactate from a 300μl of urine, as per the manufacturer's instructions. Urine creatinine was quantified using the colorimetric Creatinine Assay Kit (ab204537, Abcam plc) as per the manufacturer's instructions.

**Measurement of Urinary Metabolites**

Sample preparation

Indoxyl sulfate (M-H, *m/z* 212.0018), p-cresyl sulfate (M-H, *m/z* 187.0065), p-cresyl glucuronide (M-H, *m/z* 283.0818), hippuric acid (M-H, *m/z* 178.0504), betaine (M+H, *m/z* 118.0868), choline (M+H, *m/z* 105.1154), carnitine (M+H, *m/z* 162.1130), and nicotinamide (M+H, *m/z* 123.0558) were quantified using Ultra Performance Liquid Chromatography (UPLC) coupled to Quadrupole Time of Flight (QToF) mass spectrometry. Urine samples were prepared by addition of ice-cold acetonitrile (3:1 acetonitrile to urine) to precipitate protein, followed by incubation at -20 °C for 20 minutes and centrifugation at 20,800 x g for 10 minutes. Acetonitrile contained chlorpropamide (4 uM) and atenolol-d7 (300 ng/mL) as internal standards. To keep analytes in the linear range of the standard curve, the supernatant from urine samples was diluted with milliQ water 40-fold for indoxyl sulfate and p-cresyl sulfate, 160-fold for p-cresyl glucuronide and hippuric acid, and 5-fold for betaine. The supernatant was diluted 5-fold with acetonitrile for choline.

Chromatography and Mass Spectrometry

Indoxyl sulfate, p-cresyl sulfate, p-cresyl glucuronide, hippuric acid, betaine, carnitine and nicotinamide were separated using a Waters Acquity UPLC HSS T3 column (100 mm × 2.1 mm, 1.8 µm particle size) in a Waters Acquity UPLC I-Class system (Waters, Milford, MA). Injection volumes ranged from 0.5–2 µL between analytes. The mobile phase consisted of water + 0.1% formic acid (A) and acetonitrile + 0.1% formic acid (B) set to a flow rate of 0.45 mL/min. The UPLC gradient was as follows: 0–2 minutes 1%–60% B; 2–2.5 minutes 60% B; 2.5–3.5 minutes 80% B; 3.5–4.5 minutes 1% B. Mass spectrometry was performed using a Waters Xevo™ G2S-QToF mass spectrometer in negative (indoxyl sulfate, p-cresyl sulfate, p-cresyl glucuronide, hippuric acid) and positive (betaine, carnitine, nicotinamide) ESI modes with the following parameters: capillary voltage, 2 kV; cone voltage, 40 V; source temperature, 150 °C; desolvation temperature, 500 °C; desolvation gas flow, 1000 L/h; cone gas flow, 50 L/h. Choline was separated using a Waters Acquity BEH Amide column (100 mm × 2.1 mm, 1.7 µm particle size). The mobile phase consisted of 5 mM ammonium formate pH 3.5 (A) and acetonitrile (B) set to a flow rate of 0.45 mL/min. The UPLC gradient was as follows: 0–0.5 minutes 85% B; 0.5–1.5 minutes 85%–40% B; 1.5–2.5 minutes 40% B; 2.5–4.0 minutes 85% B. Mass spectrometry was performed in positive ESI mode with the following parameters: capillary voltage, 0.5 kV; cone voltage, 20 V; source temperature, 120 °C; desolvation temperature, 350 °C; desolvation gas flow, 1200 L/h; cone gas flow, 175 L/h. Data were acquired in sensitivity mode with a 0.05 second scan time in a 50–1200 *m/z* range and the *m/z* of each analyte was specifically targeted. Mass accuracy was maintained using a lockspray of leucine-enkephalin (1 ng/μL) measured every 10 seconds with a scan time of 0.3 seconds and averaged over 3 scans.

Quantification

Analytes were quantified using TargetLynx V4.1 software (Waters) by comparing sample peaks to a twelve-point standard curve of indoxyl sulfate (0–1200 μM), p-cresyl sulfate (0–300 μM), p-cresyl glucuronide (0–2000 μM), hippuric acid (0–8000 μM), betaine (0–700 μM), choline (0–700 μM), carnitine (0-300 μM) and nicotinamide (0-300 μM). Quality control samples contained known concentrations of each analyte, were prepared using the same protocol as biological samples and injected every nine samples. The coefficient of variation of the assay was less than 10% for all analytes.

#### Supplemental Figure Legends

**Supplemental Figure 1:** Histogram depicting the distribution of protein abundance with the QRILC imputed values. The imputed values are shown in light grey, while the measured ones are shown in dark grey. As expected, the imputed values represent low abundance proteins.

**Supplemental Figure 2**: Heatmap generated using unsupervised hierarchical clustering depicting the expression profiles of the 70 DE proteins using the non-imputed data (with missing values assumed to be "0"). Row annotations highlight distinct clusters of proteins exhibiting co-expression across samples. Annotation of the columns details the experimental group and timepoint.

**Supplemental Figure 3**: Full western blot images. Western blot images were acquired using the DNR Bioimaging Systems MicroChemi 4.2. which incorporates the molecular weight reference image with the image of the membrane.

**Supplemental Figure 4**: Relative mRNA expression of PPAR-family and related transcription factors at baseline. Significance assessed by Mann Whitney test, n=3-5 per group.

**Supplemental Figure 5: Analysis of Urine Metabolites**

The absolute urinary concentration of **(A)** Indoxyl Sulfate, Hippuric Acid, p-Cresyl Glucuronide, p-Cresyl Sulfate, and **(B)** Betaine and Choline were measured (μmol/L) and normalized to urinary creatinine (μmol/L). Values at POD3 are expressed as fold change over the baseline value. N= 5 urines per group. Urinary concentration of **(C)** Glucose and **(D)** Lactate were measured (both mmol/L) at POD3, and normalised to urinary creatinine (μmol/L). N=3-4 urines per group. Differences between groups were assessed by Mann-Whitney test, *$p<0.05$ compared to SCS at POD3. NEVKP, normothermic ex vivo kidney perfusion; POD3, post-operative day 3; SCS, static cold storage.

#### Supplemental Figures 1-5

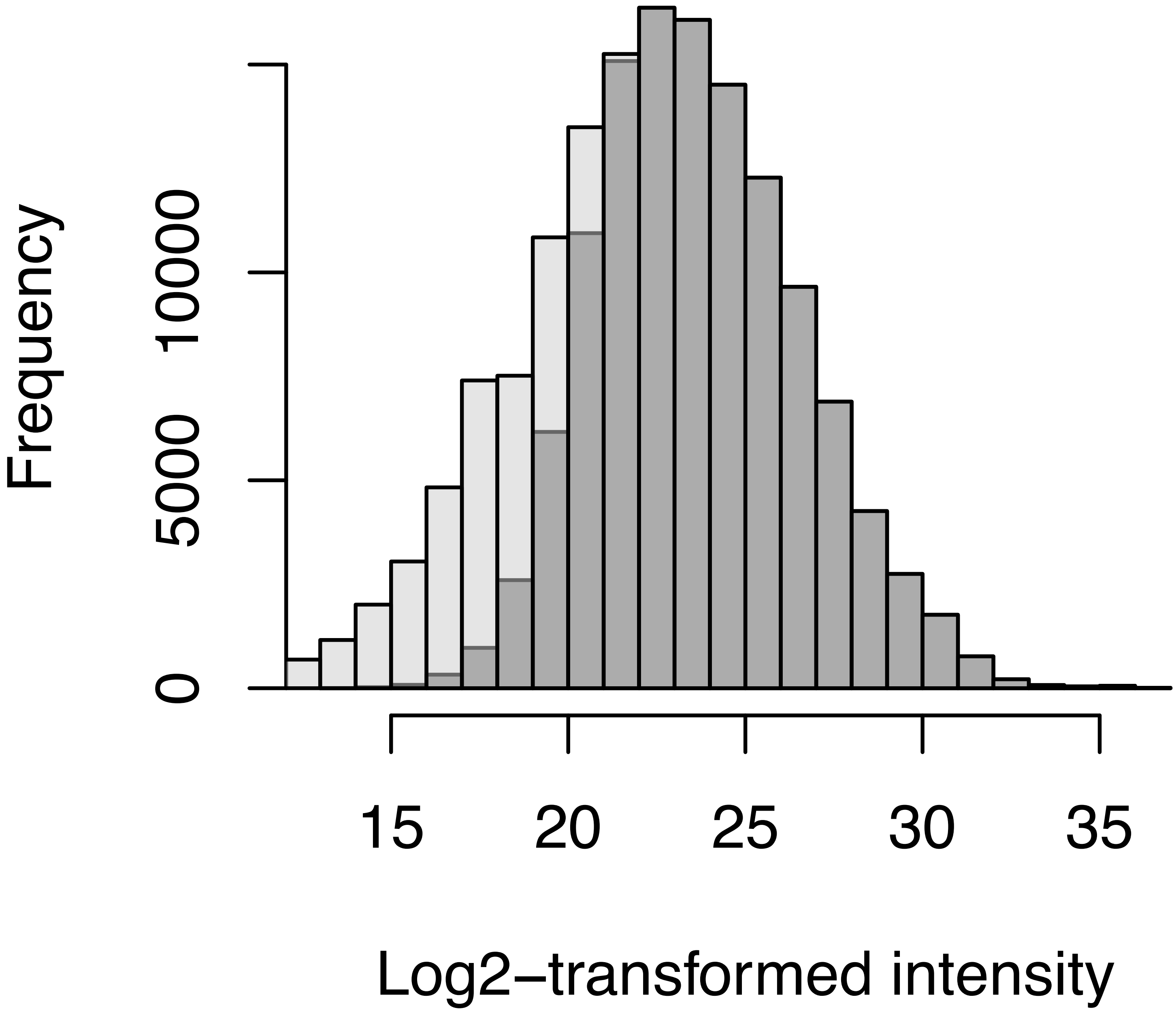

Frequency
0
5000
10000
15
20
25
30
35
Log2−transformed intensity

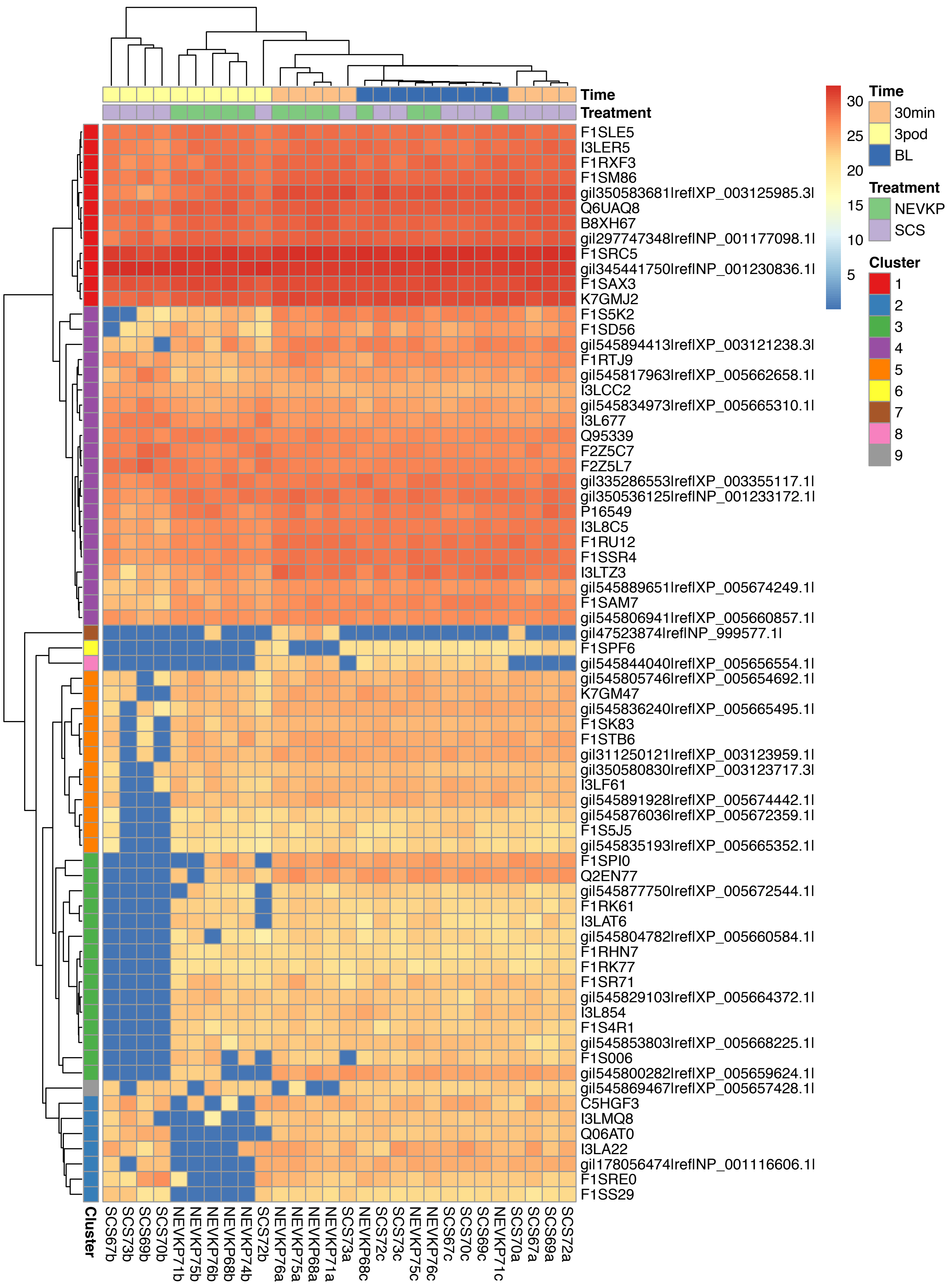

Time
Treatment
F1SLE5
I3LER5
F1RXF3
F1SM86
gi|350583681|ref|XP_003125985.3|
Q6UAQ8
B8XH67
gi|297747348|ref|NP_001177098.1|
F1SRC5
gi|345441750|ref|NP_001230836.1|
F1SAX3
K7GMJ2
F1S5K2
F1SD56
gi|545894413|ref|XP_003121238.3|
F1RTJ9
gi|545817963|ref|XP_005662658.1|
I3LCC2
gi|545834973|ref|XP_005665310.1|
I3L677
Q95339
F2Z5C7
F2Z5L7
gi|335286553|ref|XP_003355117.1|
gi|350536125|ref|NP_001233172.1|
P16549
I3L8C5
F1RU12
F1SSR4
I3LTZ3
gi|545889651|ref|XP_005674249.1|
F1SAM7
gi|545806941|ref|XP_005660857.1|
gi|47523874|ref|NP_999577.1|
F1SPF6
gi|545844040|ref|XP_005656554.1|
gi|545805746|ref|XP_005654692.1|
K7GM47
gi|545836240|ref|XP_005665495.1|
F1SK83
F1STB6
gi|311250121|ref|XP_003123959.1|
gi|350580830|ref|XP_003123717.3|
I3LF61
gi|545891928|ref|XP_005674442.1|
gi|545876036|ref|XP_005672359.1|
F1S5J5
gi|545835193|ref|XP_005665352.1|
F1SPI0
Q2EN77
gi|545877750|ref|XP_005672544.1|
F1RK61
I3LAT6
gi|545804782|ref|XP_005660584.1|
F1RHN7
F1RK77
F1SR71
gi|545829103|ref|XP_005664372.1|
I3L854
F1S4R1
gi|545853803|ref|XP_005668225.1|
F1S006
gi|545800282|ref|XP_005659624.1|
gi|545869467|ref|XP_005657428.1|
C5HGF3
I3LMQ8
Q06AT0
I3LA22
gi|178056474|ref|NP_001116606.1|
F1SRE0
F1SS29
Cluster
SCS67b
SCS73b
SCS69b
SCS70b
NEVKP71b
NEVKP75b
NEVKP76b
NEVKP68b
NEVKP74b
SCS72b
NEVKP76a
NEVKP75a
NEVKP68a
NEVKP71a
SCS73a
NEVKP68c
SCS72c
SCS73c
NEVKP75c
NEVKP76c
SCS67c
SCS70c
SCS69c
NEVKP71c
SCS70a
SCS67a
SCS69a
SCS72a
30
25
20
15
10
5
Time
30min
3pod
BL
Treatment
NEVKP
SCS
Cluster
1
2
3
4
5
6
7
8
9

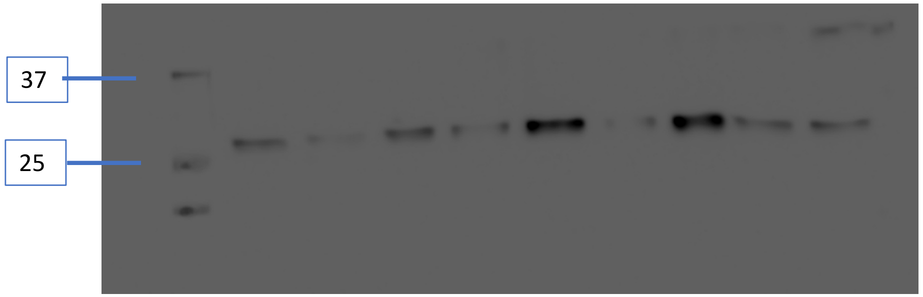


ETFB

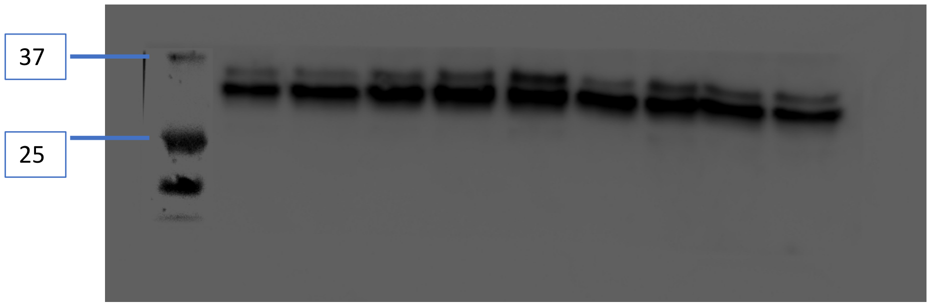


Blot reprobed for GAPDH

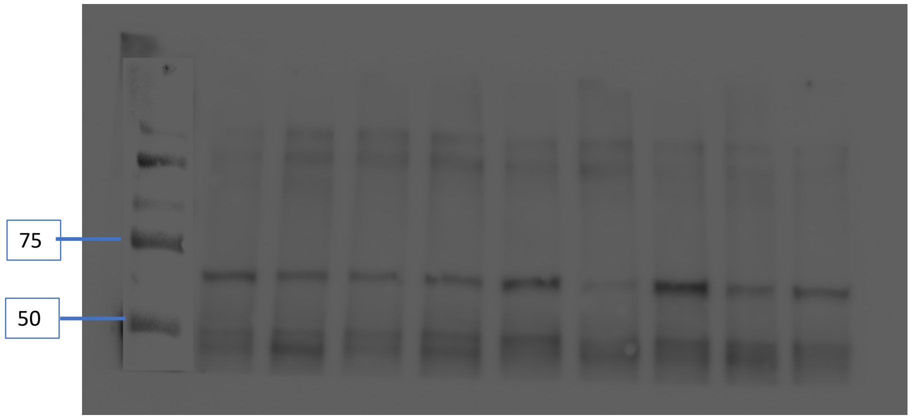


Blot transected, top probed for CPT2

Bottom probed for GAPDH

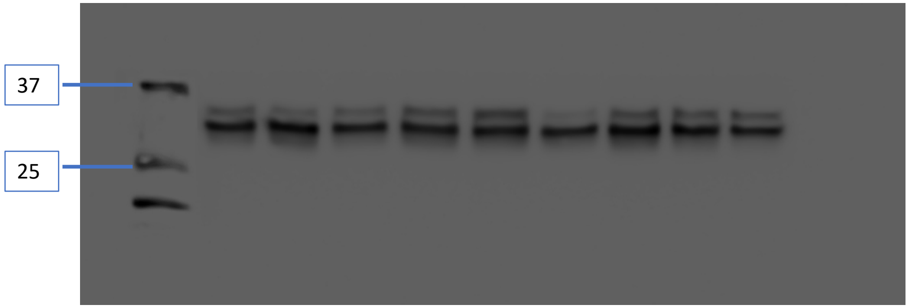

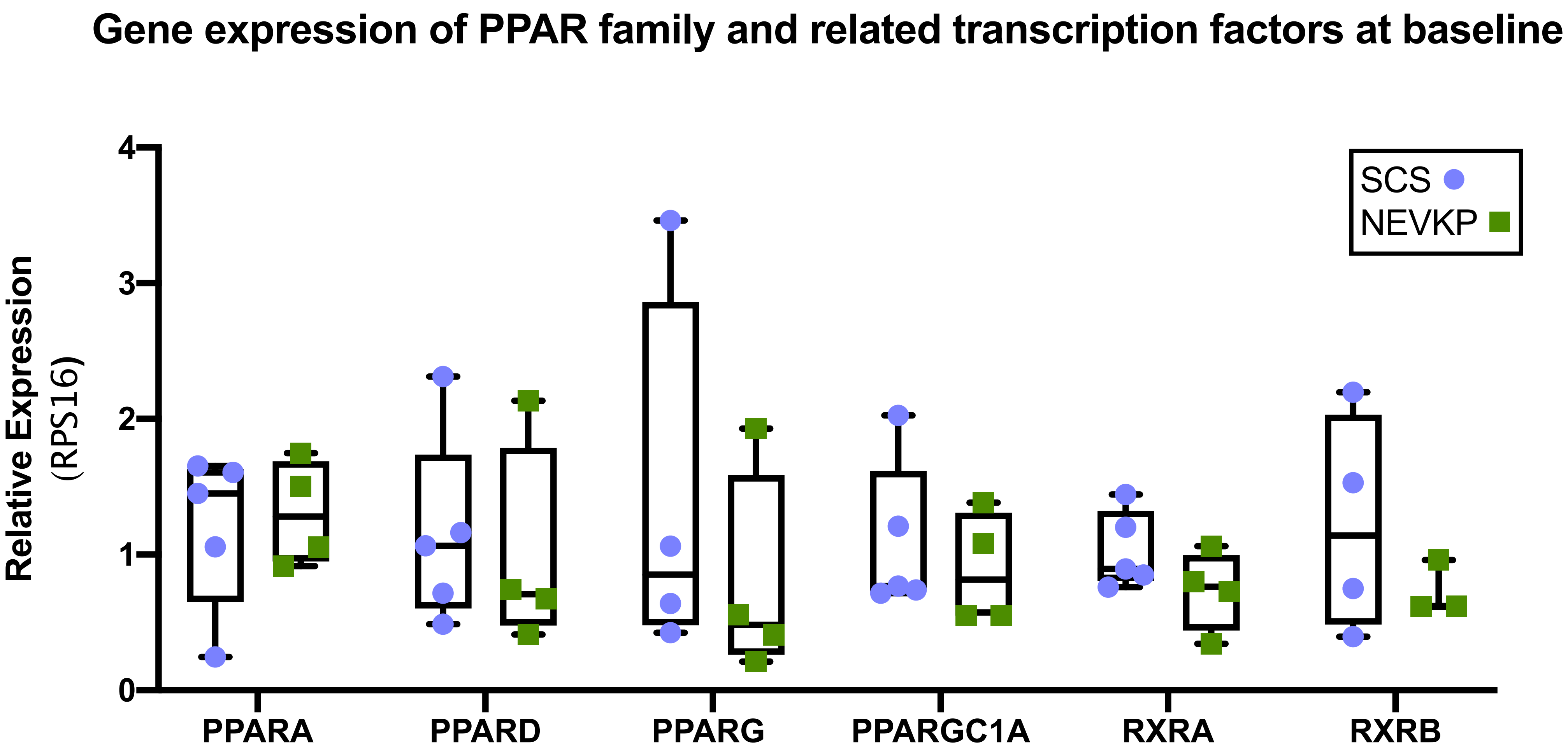

Gene expression of PPAR family and related transcription factors at baseline
Relative Expression
(RPS16)
0
1
2
3
4
PPARA
PPARD
PPARG
PPARGC1A
RXRA
RXRB
SCS
NEVKP

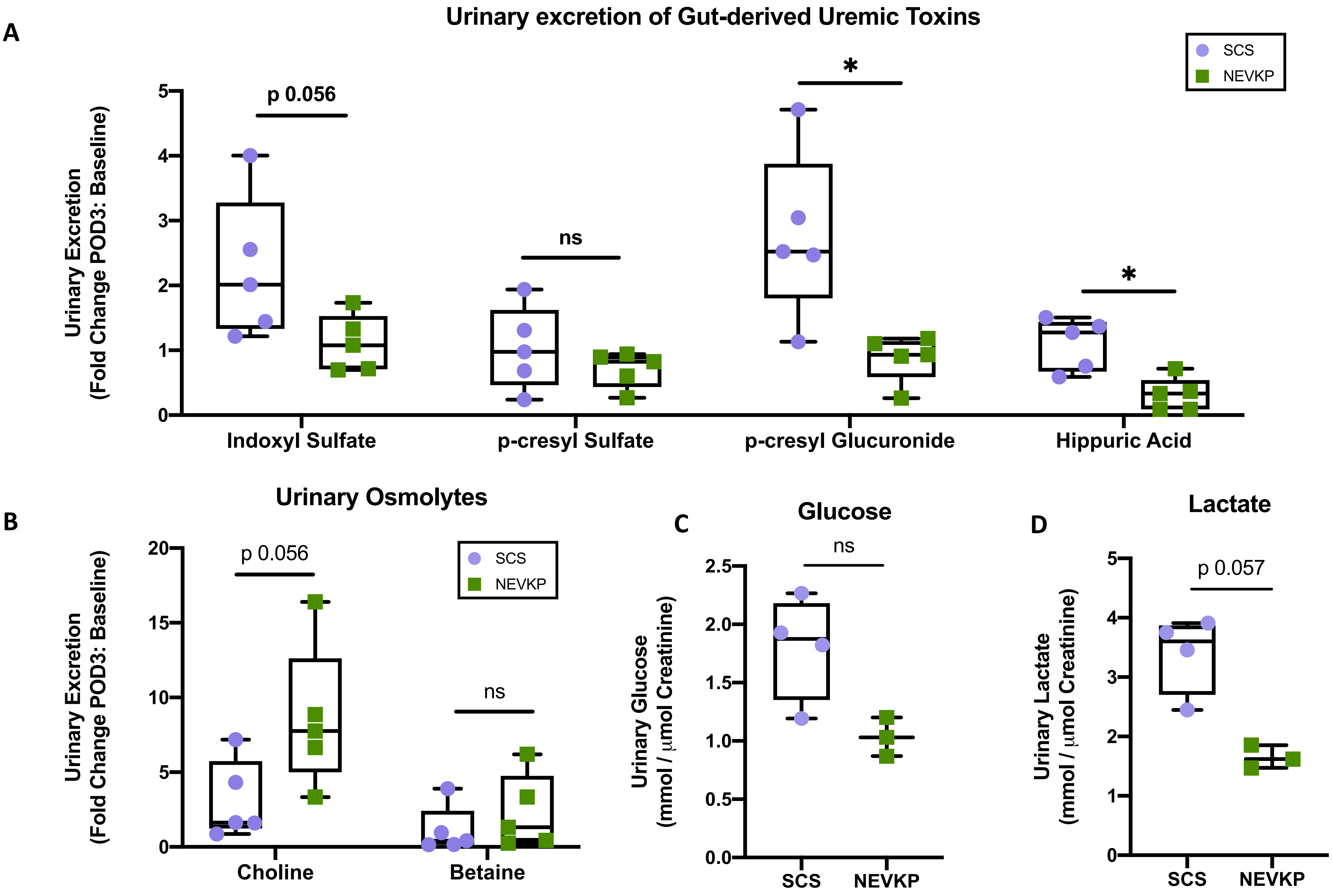
A
Urinary excretion of Gut-derived Uremic Toxins
SCS
NEVKP
Urinary Excretion
(Fold Change POD3: Baseline)
p 0.056
ns
*
*
Indoxyl Sulfate
p-cresyl Sulfate
p-cresyl Glucuronide
Hippuric Acid
B
Urinary Osmolytes
p 0.056
ns
Choline
Betaine
C
Glucose
Urinary Glucose
(mmol / μmol Creatinine)
ns
SCS
NEVKP
D
Lactate
Urinary Lactate
(mmol / μmol Creatinine)
p 0.057
